# Galectin-1 induces macrophage immunometabolic reprogramming, modulates T cell immunity and attenuates atherosclerotic plaque formation

**DOI:** 10.1101/2025.11.21.689721

**Authors:** Ya Li, Julian Leberzammer, Xavier Blanchet, Rundan Duan, Michael Lacy, Vasiliki Triantafyllidou, Veit Eckardt, Eva Briem, Anna Simone Jung, Rui Su, Joel Guerra, Yvonne Jansen, Michael Hristov, Wolfgang Enard, Jürgen Bernhagen, Christian Weber, Dorothee Atzler, Alexander Bartelt, Yvonne Doring, Donato Santovito, Herbert Kaltner, Anna-Kristin Ludwig, Philipp von Hundelshausen

## Abstract

**Background and aims:** Atherosclerosis is a chronic immunometabolic disease driven by lipid accumulation and immune cell infiltration. Macrophages and T cells play key roles throughout plaque development. Galectin-1 (Gal-1), a glycan-binding protein, modulates immune functions in these cells and has been reported to attenuate atherosclerosis, though its mechanisms remain incompletely understood. Here, we investigated the effects of Gal-1 on macrophages and T cells during plaque formation.

**Methods:** Effects of Gal-1 on atherosclerosis, macrophages and T cells during lesion formation were studied in *Apoe*^−/−^ mice treated with recombinant Gal-1. Complementary mouse peritoneal foam cell and in vitro macrophage and T cell cultures experiments were performed to study T cell differentiation, macrophage function, polarization end energy metabolism. The impact of Gal-1 on human macrophages was further evaluated in endarterectomy specimens.

**Results:** Gal-1 treatment reduced lesion size and increased circulating IL-10 levels, inversely correlating with plaque burden. Unexpectedly, IL-10 neutralization also mitigated atherosclerosis, indicating that its action is at least partially IL-10–independent. In plaques, Gal-1 promoted anti-inflammatory macrophage phenotypes, mirrored by a quiescent metabolic and anti-inflammatory profile in foamy macrophages ex vivo. The use of the Gal-1^E71Q^ variant revealed that these effects were only partly dependent on glycan binding. Beyond IL-10, Gal-1 reshaped cytokine profiles by increasing IL-17, IL-22, and IL-23, consistent with a macrophage-driven regulatory Th17 response, alongside higher frequencies of IL-10–producing and regulatory T cells.

**Conclusion:** Gal-1 protects against atherosclerosis associated with reprogramming macrophages and tuning T cell immunity through glycan-dependent and –independent pathways.

## 1. Introduction

Atherosclerosis, a chronic immunometabolic disease of the arterial wall, is the primary underlying cause of cardiovascular disease (CVD), which remains the leading cause of global death. Despite improvements in life expectancy, in low– and middle-income countries where the prevalence of CVD is expected to rise [1]. Central to the development and progression of atherosclerosis is the interplay between dyslipidemia and chronic inflammation, particularly the activation and infiltration of immune cells into the intimal layer of the arterial wall [2]. Macrophages and T cells, in particular, are pivotal players in plaque formation and contribute to the maladaptive immune response that drives disease progression [3].

Traditional therapies for atherosclerosis focus on lipid-lowering and thrombosis prevention, but recent clinical trials—such as the Canakinumab Anti-inflammatory Thrombosis Outcomes Study (CANTOS), the Colchicine Cardiovascular Outcomes Trial (COLCOT), and Low Dose Colchicine (LoDoCo)—have highlighted the therapeutic potential of targeting inflammation to reduce cardiovascular events [4]. However, the cost-benefit ratio of these approaches and substantial side effects have hindered their widespread clinical adoption, underscoring the need for novel therapeutic strategies.

Among potential anti-inflammatory molecules, Gal-1, an endogenous glycan-binding protein of the galectin family, has emerged as a promising candidate with immunomodulatory effects on various immune cells, including macrophages, T cells, neutrophils and B cells [5–7]. By binding β-galactoside units on glycoproteins on cell surfaces and the extracellular matrix, Gal-1 regulates cell migration, activation, and apoptosis, thereby playing a critical role in all stages of inflammation—from initiation to resolution. Its bi– or multivalent nature enables cis or trans crosslinking of binding partners, further influencing these processes [8]. Beyond glycan binding, galectin-1 and –3 also interact with chemokines and intracellular transcription factors, such as forkhead box P3 (Foxp3), altering leukocyte migration and gene expression [9, 10].

Modulation of monocyte/macrophage polarization towards an anti-inflammatory and pro-resolving phenotype has been reported [11]. Additionally, Gal-1 has been shown to inhibit leukocyte-endothelial interactions, further promoting an anti-inflammatory phenotype [12]. Notably, both pro– and anti-inflammatory effects of Gal-1 exist depending on tissue context, intracellular or extracellular localization, pathologic conditions, and spatiotemporal expression of other regulatory programs [13].

Gal-1 has emerged as an important mediator of cardiovascular inflammation [14]. Circulating levels of Gal-1 are elevated following atherosclerotic events such as large artery stroke, and mouse models suggest that Gal-1 promotes post-stroke healing through interactions with neurons [15]. Moreover, Gal-1-deficient mice exhibit exacerbated cardiac inflammation and worsened vascular remodeling following myocardial infarction. It has been recently reported that Gal-1-deficient mice present with an increased atherosclerotic plaque burden and that exogenous Gal-1 attenuates atherosclerotic lesion development via effects on smooth muscle cells [16, 17].

Given the critical roles of macrophages and T cells in atherosclerosis, and the emerging evidence that Gal-1 affects their biology, we sought to additionally investigate how Gal-1 influences immunometabolic reprogramming of macrophages and the immunity of T cells in the context of atherosclerosis.

## 2. Methods and materials

Comprehensive methods are available in the **Supplementary Information online**.

### 2.1 Animals and procedures

8-week-old *Apoe*^−/−^ mice (C57BL/6J background, Charles River Laboratories, France) were fed a Western diet (WD; 21% fat, 0.2% cholesterol, Ssniff, TD88137) for 8 weeks. They received intraperitoneal (i.p.) injections of either human recombinant Gal-1 (100 µg in 100 µL PBS, three times weekly) or PBS, as established previously [16]. In one setup, mice also received neutralizing IL-10 Ab (InVivoMAb, JES5-2A5, 500 µg i.p., three times a week during the first week and 300 µg twice a week thereafter), while controls received PBS or IL-10 Ab alone [18]). Feces were collected before sacrifice. Mice were euthanized under anesthesia (30 mg/kg xylazine, 150 mg/kg ketamine) following final blood sampling with subsequent perfusion and organ collection.

Immune cell and peritoneal foamy macrophage accumulation were assessed in additional experiments. Female *Apoe*^−/−^ mice were fed a WD for 4 weeks. Starting in week 3, mice received Gal-1 or Gal-1^E71Q^ (100 µg, three times weekly i.p.) or PBS. After the dietary run-in phase, mice were injected with 750 µL of 4% thioglycolate. Four days post-injection, mice received a final injection of Gal-1, Gal-1^E71Q^, or PBS, followed by an 8-hour fasting period. Peritoneal lavage fluid was collected by washing the abdominal cavity with 10 mL ice-cold PBS containing 0.1% BSA and 2 mM EDTA. In one setup, female *Apoe*^−/−^ mice were fed a WD for 4 weeks, and then mice were sacrificed and peritoneal lavage was collected.

All experiments were approved by local authorities (Regierung von Oberbayern, Germany; ROB-55.2Vet-2532.Vet_02-17-180, ROB-55.2-2532.Vet_02-22-150) and in compliance with the guidelines of Directive 2010/63/EU of the European Parliament on the protection of animals used for scientific purposes.

### 2.2 Expression and purification of Gal-1 and Gal-1^E71Q^

Recombinant galectins were generated using pGEMEX-1 expression plasmid and the pGEX-6p2 as described [19]. A detailed protocol can be found in supplementary Information online.

### 2.3 Peritoneal macrophage experiment

Peritoneal lavage was collected from mice fed a WD as described above. Erythrocytes were lysed using ACK buffer. Peritoneal cells (1 × 10^6^ cells/well) were seeded onto 12-well cell culture plates and incubated with or without 100 nM Gal-1 for 24 hours at 37 °C. Cells were then collected and polarization of peritoneal macrophages (CD206, CD80, CD86, MHC-II) were assessed by flow cytometry.

### 2.4 Extracellular Flux X24 Seahorse Measurement

Oxygen consumption rate (OCR) and extracellular acidification rate (ECAR) were measured on an XF-24 Extracellular Flux Analyzer (Agilent) (**detailed in Supplementary Information online**). Briefly, BMDMs or peritoneal macrophages (1 × 10^5^ cells/well) were seeded onto seahorse cell culture plates. Measurements were performed in triplicate. The following compounds were used: oligomycin (1.5 µM), fluoro carbonyl cyanide phenylhydrazone (FCCP, 1.5 µM), and rotenone + antimycin A (Rtn/AA, 2.5 µM) for the Mito Stress Test; glucose (10 mM), oligomycin (1.5 µM), and 2-deoxy-D-glucose (2-DG, 50 µM) for the Glycolysis Stress Test. After the readout, cells were lysed, and protein concentrations were determined using the BCA Protein Assay Kit (Thermo Scientific™ 23225). All metabolic rates were normalized to total protein levels.

### 2.5 RNA sequencing and analysis

RNA-sequencing was performed from inflammatory peritoneal macrophages using the prime-seq protocol [20]. Details on protocol and analysis can be found in the **Supplementary Information, Methods.**

### 2.6 Cholesterol and triglycerides measurement

Total cholesterol and triglycerides concentrations were determined using the enzymatic colorimetric method (Cholesterol-CHOD-PAP, Triglycerides-GPO-PAP, Roche Diagnostics) according to the manufacturer’s instructions. Lipids were extracted from 100 mg of liver and feces with a chloroform/methanol solution, and dried pellets were dissolved in buffer (0.1 M potassium phosphate, pH 7.4, 0.05 M NaCl, 5 mM cholic acid, 0.1% Triton X-100).

### 2.7 Statistical analysis

Statistical analyses were performed using GraphPad Prism version 10 (GraphPad Software, LLC). Data distribution was assessed for normality using the Shapiro-Wilk test, and lognormality was tested via logarithmic transformation. If the data were normally distributed or could be transformed into a normal distribution, parametric tests (paired or unpaired Student’s t-test, or ANOVA for comparisons of more than two groups) were applied. For non-normally distributed data, the non-parametric Mann-Whitney test was used. One-way ANOVA was followed by Holm-Šídák’s post hoc test for group comparisons. A two-tailed *p* value of < 0.05 was considered statistically significant. Data are presented as mean ± standard error of the mean (SEM), unless otherwise specified.

## 3. Results

### 3.1 Administration of Gal-1 reduces atherosclerotic plaque area and plasma cholesterol levels

To assess the impact of Gal-1 treatment on atherosclerotic lesions, 8-week-old *Apoe*^−/−^ mice were fed a WD for 8 weeks while receiving three times weekly intraperitoneal injections of either 100 µg of human recombinant Gal-1 or PBS (**Fig.1A**). Histological analysis of serial sections from the aortic root, spanning from the proximal to the distal regions, revealed a significant reduction in the absolute plaque area by approximately 40% in Gal-1-treated mice compared to PBS-treated controls (**Fig.1B**), well in line with a previously observed atheroprotective effect of Gal-1 in another model of atherosclerosis [16]. Similarly, the relative plaque area, calculated as the percentage of total plaque area within the three aortic leaflets compared to the total root area, was also significantly reduced in Gal-1-treated mice (**Fig.1C**). Gal-1 did not show any sex-related differences in the plaque reduction; therefore, only female mice were used in the subsequent experiments. Necrotic cores, defined as hypocellular or acellular regions within the plaque, were generally small at this early stage of atherosclerosis. Neither the size of the necrotic core, nor the thickness of the fibrous cap, the eosinophilic layer overlaying the necrotic core, were altered across groups (**Fig.1D**). Furthermore, collagen content of the plaques revealed no changes following the Gal-1 treatment (**Fig.1E**). These findings indicate that Gal-1 treatment reduces atherosclerotic plaque size but does not alter plaque stability.

**Fig. 1.**
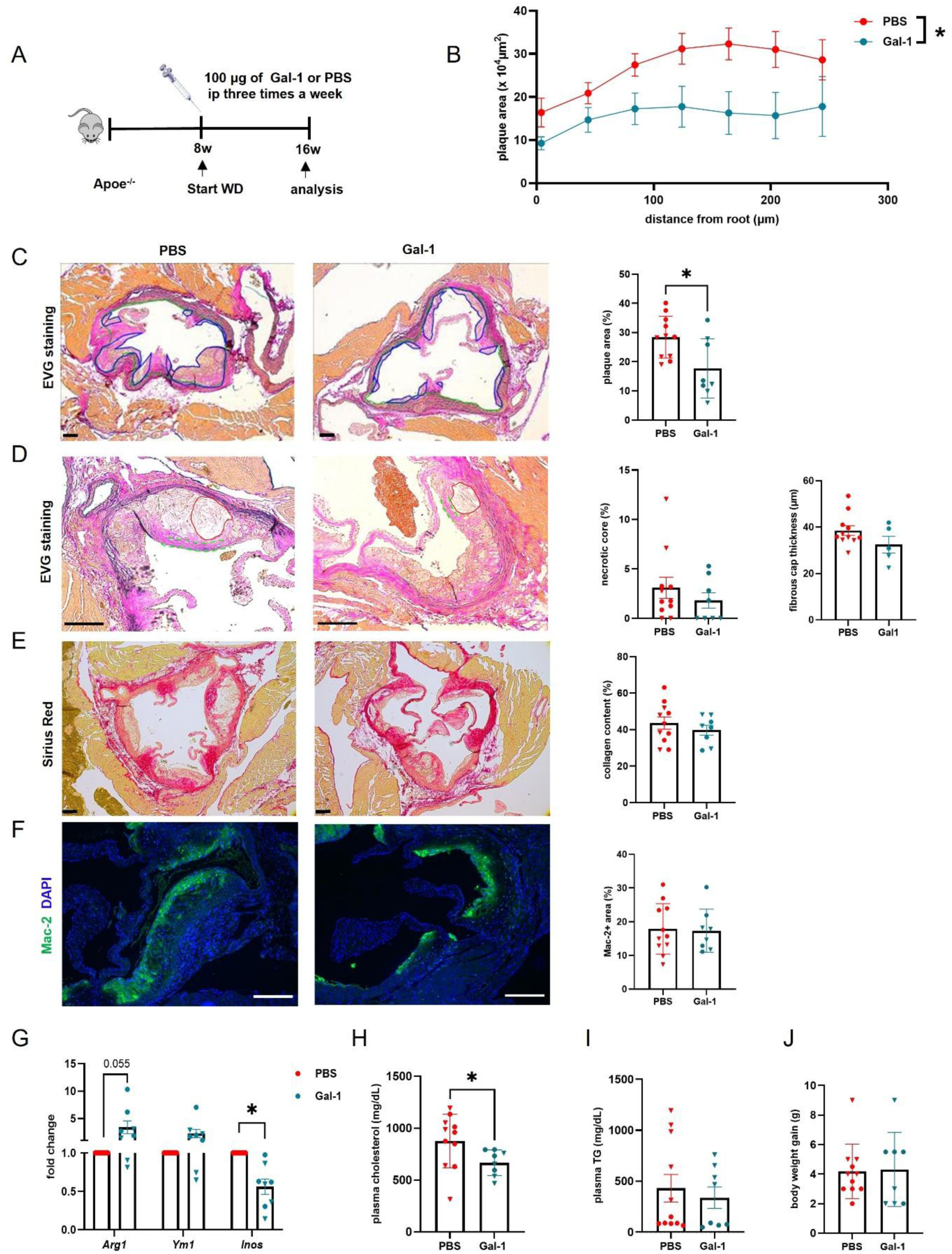
Administration of Gal-1 reduces atherosclerotic plaque formation and plasma cholesterol levels. **(A)** Experimental design. (**B)** Displayed is the plaque area from the proximal to distal part of the aortic roots from PBS-(n=11, five females) and Gal-1-treated (n=8, four females) *Apoe*^−/−^ mice after 8 weeks on a Western diet (WD). The area under the curve was used to compare overall plaque burden. **(C)** Representative images of aortic roots stained with elastin/van Gieson (EVG) showing plaque boundaries (blue outline) and quantification of relative plaque area. **(D)** Representative images of aortic roots stained with EVG showing necrotic core (red outline), fibrous cap thickness measurements (dashed green outline), and their quantification. **(E)** Representative images of Sirius Red staining and quantification of collagen deposition in aortic root plaques. **(F)** Representative immunofluorescence images and quantification of macrophage (Mac-2⁺) infiltration relative to plaque area. **(G)** Expression of inflammatory genes related to macrophage profiling in the aortic arch of PBS-(n=8) and Gal-1-treated mice (n=8) after 8 weeks of WD. **(H)** Plasma cholesterol and **(I)** triglyceride (TG) levels. **(J)** Body weight gain after 8 weeks of WD. Data are expressed as mean ± SEM, with individual data points shown. Bullets represent female mice, and triangles represent male mice. \**p* < 0.05, two-tailed Student’s t-test. Scale bar: 100 μm.

In terms of cellular composition, the area of lesional macrophages remained unchanged (**Fig.1F**). Absolute CD3-positive T cells were reduced in the plaques of Gal-1-treated mice, while their frequency relative to plaque area remained unaltered (**Fig.S1A-C**). To further characterize the plaque phenotype, the inflammatory status of the aortic arch was assessed. The anti-inflammatory marker *Arg1* tended to be upregulated, and pro-inflammatory *Inos* was significantly decreased in Gal-1-treated mice compared to PBS-injected mice, indicating a shift toward an anti-inflammatory milieu (**Fig.1G**). These findings suggest that Gal-1 attenuates plaque development and promotes a less inflammatory plaque environment.

Consistent with a previous study showing that genetic deletion of *Lgals1,* the gene encoding Gal-1, results in elevated blood cholesterol levels, we observed a significant 24% reduction in plasma cholesterol levels in Gal-1-treated mice (**Fig.1H**) [16]. However, no significant differences were detected in triglyceride levels, body weight gain or body weight (**Fig.1I-J, Fig.S1D**).

### 3.2 Additive effects of IL-10 neutralization and Gal-1 in reducing atherosclerosis

Consistent with the anti-inflammatory effects of Gal-1, we observed a significant increase in the potent anti-inflammatory cytokine IL-10 in the plasma of Gal-1-treated mice (**Fig.2A**). Notably, plasma IL-10 levels negatively correlated with plaque size, with higher IL-10 levels being associated with smaller plaques (**Fig.2B**). In line with this finding, Gal-1 has been reported to mediate immunomodulatory effects through IL-10, and IL-10 injections have also been reported to lower cholesterol levels in humans [18, 21]. To determine whether blocking circulating IL-10 would negate the beneficial effects of Gal-1 on atherosclerosis and cholesterol levels, we administered a neutralizing IL-10 antibody, using a previously established experimental protocol, via intraperitoneal injection, combined with the Gal-1 treatment regimen for 4 weeks (**Fig.2C**) [18].

**Fig. 2.**
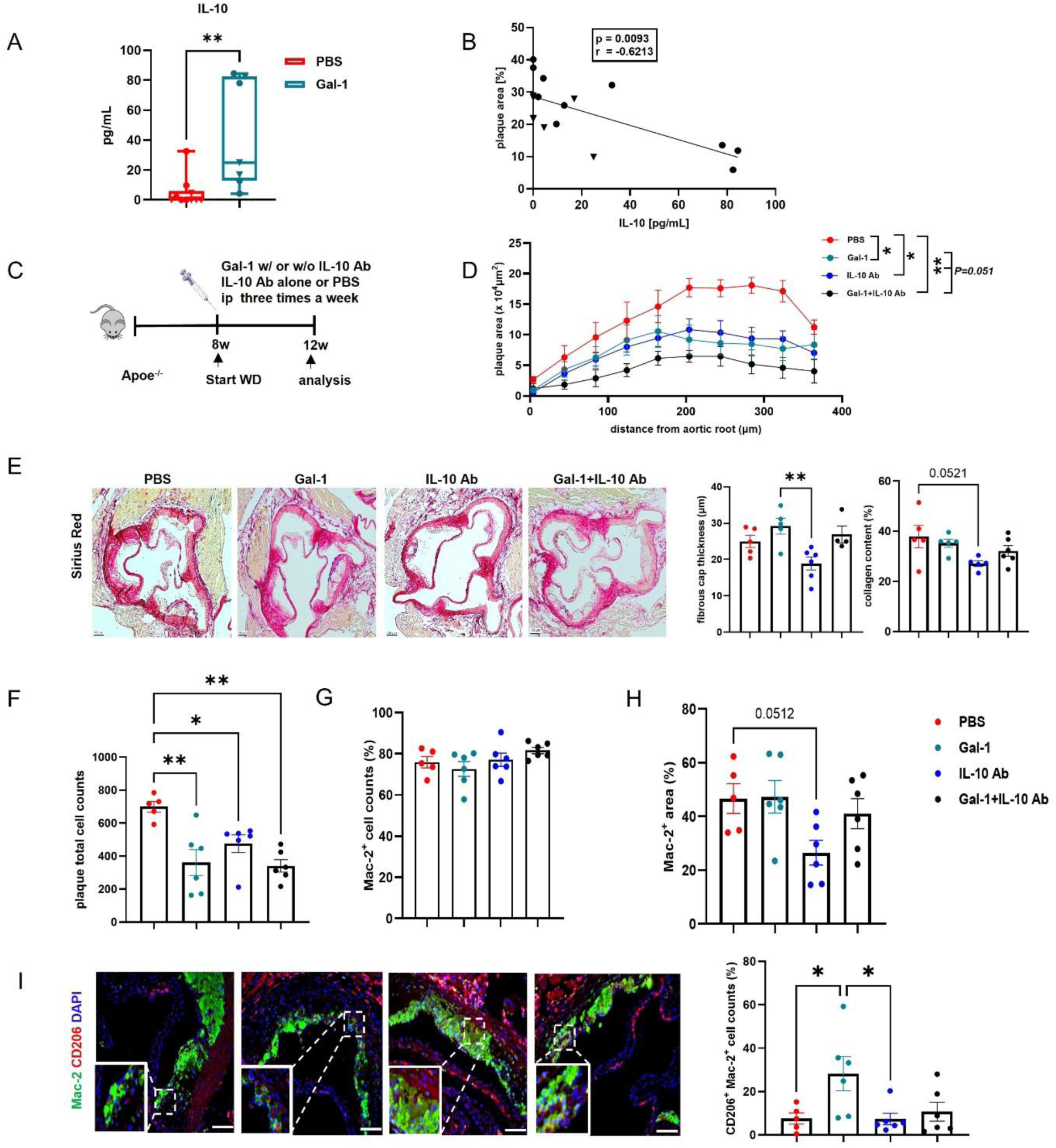
Additive effects of IL-10 neutralization and Gal-1 in reducing atherosclerosis. **(A)** Plasma IL-10 levels from PBS-(n=10) and Gal-1-treated *Apoe*^−/−^ mice (n=7) after 8 weeks of WD as shown in **Fig.1A**. **(B)** Spearman’s correlation between plasma IL-10 levels and aortic root plaque area**. (C)** Experimental design. Female *Apoe*^−/−^ mice were treated with PBS, Gal-1, IL-10 Ab (neutralizing IL-10 antibody, clone JES5-2A5), or Gal-1 + IL-10 Ab following 4 weeks of WD (PBS: n=5, other groups: n=6). **(D)** Displayed is the plaque area from the proximal to distal part of the aortic roots resulting in the area under the curve. **(E)** Stability of the plaque was assessed in Sirius Red-stained aortic root sections showing representative images, fibrous cap (FC) thickness measurements (dashed blue outline) and quantification of collagen content. FC was not quantifiable for two mice in the Gal-1 + IL-10 Ab group. **(F)** Plaque cellularity was quantified by counting DAPI-positive nuclei**. (G-H)** Percentage of macrophage numbers (DAPI^+^Mac-2^+^, **G**) and Mac-2^+^-area **(H)** is indicated per plaque area. **(I)** Representative immunofluorescence images and quantification of numbers of CD206^+^Mac-2^+^ macrophages relative to total plaque cells. **(A) is expressed as box plots showing the interquartile range, median, and minimum to maximum, analyzed by Mann-Whitney U-test. (A, B)** Bullets represent female mice and triangles represent male mice. **(D-I)** are presented as mean ± SEM and analyzed by **One-**way ANOVA followed by Holm-Šídák’s post hoc test. \**p* < 0.05 and \*\**p* < 0.01. Scale bar: 100 μm.

After 4 weeks of WD, Gal-1 treatment significantly reduced atherosclerotic lesions, primarily in the aortic root and to a lesser extent in the aortic arch. Unexpectedly, blocking IL-10 did not exacerbate plaque formation; it led to a significant reduction in plaque size, and the combination of anti-IL-10 and Gal-1 resulted in a further numerical reduction, though non-significant, compared to either treatment alone (**Fig.2D, Fig.S2A**).

There were no significant differences in the necrotic core size, fibrous cap thickness, or collagen content between PBS– and Gal-1-treated groups. In contrast, IL-10 blockade resulted in a thinner fibrous cap and reduced collagen content (p=0.05), suggesting a shift toward a more unstable plaque phenotype (**Fig.2E, Fig.S2B)**.

Gal-1 treatment reduced plaque cellularity in line with the smaller plaque size, and cellularity was likewise reduced by anti–IL-10 alone or in combination with Gal-1 (**Fig.2F**). Regarding macrophages, the proportion among total plaque cells remained similar across groups (**Fig.2G**). However, the Mac-2 (Gal-3)–positive area was reduced only by anti–IL-10 treatment associated with a decrease in circulating monocytes (**Fig.2H, Fig.S2C**). A plausible explanation for the decreased Mac-2–positive area in the anti–IL-10-treated group, despite unchanged macrophage numbers, is that anti–IL-10 treatment may have altered Gal-3 release, expression or reduced macrophage size, in line with a report showing that IL-10 can promote Gal-3 expression in a STAT3-dependent manner [22]. The number of anti-inflammatory macrophages (CD206^+^Mac-2^+^) per total plaque cells was increased following Gal-1 treatment (**Fig.2I**) indicating that Gal-1 promotes an anti-inflammatory macrophage phenotype.

T cell numbers and lipid accumulation in the plaque, measured by the intracellular lipid droplet marker perilipin-2, were similar across groups (**Fig.S2D, E**).

Taken together, these results demonstrate that Gal-1 influences atherosclerosis through mechanisms that are not solely dependent on IL-10, while IL-10 neutralization reduces plaque size and promotes features of instability.

### 3.3 Gal-1 administration and IL-10 blockade reduce plasma cholesterol without affecting intestinal lipid absorption or liver lipid metabolism

In the 4-week trial, Gal-1 and/or IL-10 antibody treatment reduced plasma cholesterol by ∼20%, consistent with the 8-week results. The combination of Gal-1 and anti-IL-10 also reduced cholesterol, but the effect was not additive (**Fig.S3A**). Given the association between IL-10 blockade and increased risk of enterocolitis and inflammatory bowel disease (IBD), we considered that the observed reduction could be attributed to decreased dietary cholesterol absorption or enhanced hepatic metabolism [23]. Throughout the trial, all mice showed similar weight gain (**Fig.S3B, C**), with no signs of diarrhea, bloody stools, or colitis. Stool cholesterol content, and triglyceride levels, were comparable across groups (**Fig.S3D-F**). Histological analysis of liver sections stained with Oil Red O (ORO) showed no lipid accumulation (**Fig.S3G, H**), and liver cholesterol and triglyceride levels were similar between groups (**Fig.S3I, J**). Additionally, hepatic expression of genes related to lipid metabolism showed no significant differences (**Fig.S3K**). Moreover, Gal-1 has been shown to bind to exosomes, and it is conceivable that it might also bind to lipoproteins, so that cholesterol depletion via an extracellular vesicle (EV)-mediated process is deemed possible [24]. Gal-1 addition to plasma caused precipitation, but the precipitate lacked detectable cholesterol and did not reduce total cholesterol levels compared to controls (**Fig.S3L).** These findings suggest that neither cholesterol resorption, altered hepatic metabolism nor binding to extracellular vesicles contribute to the reduced plasma cholesterol levels in Gal-1 treated mice.

### 3.4 Effects of Gal-1 on energy metabolism and BMDM polarization are independent of oxLDL

We next hypothesized that Gal-1 might modulate lipoprotein uptake by macrophages and thereby influence plaque inflammation. When bone marrow-derived macrophages (BMDMs) were incubated with Gal-1 and oxLDL, spontaneous uptake of oxLDL occurred after 4 and 24 hours, which was significantly enhanced by Gal-1 (**Fig.3A, B**). Stimulation with oxLDL alone and combined with Gal-1 increased the gene expression of *Cd36*, and combination with Gal-1 significantly upregulated CD36 surface expression (**Fig.3C, D**), suggesting that under conditions where LDL is oxidized, Gal-1 enhances its uptake by macrophages, which may affect foam cell formation and plaque inflammation.

**Fig. 3.**
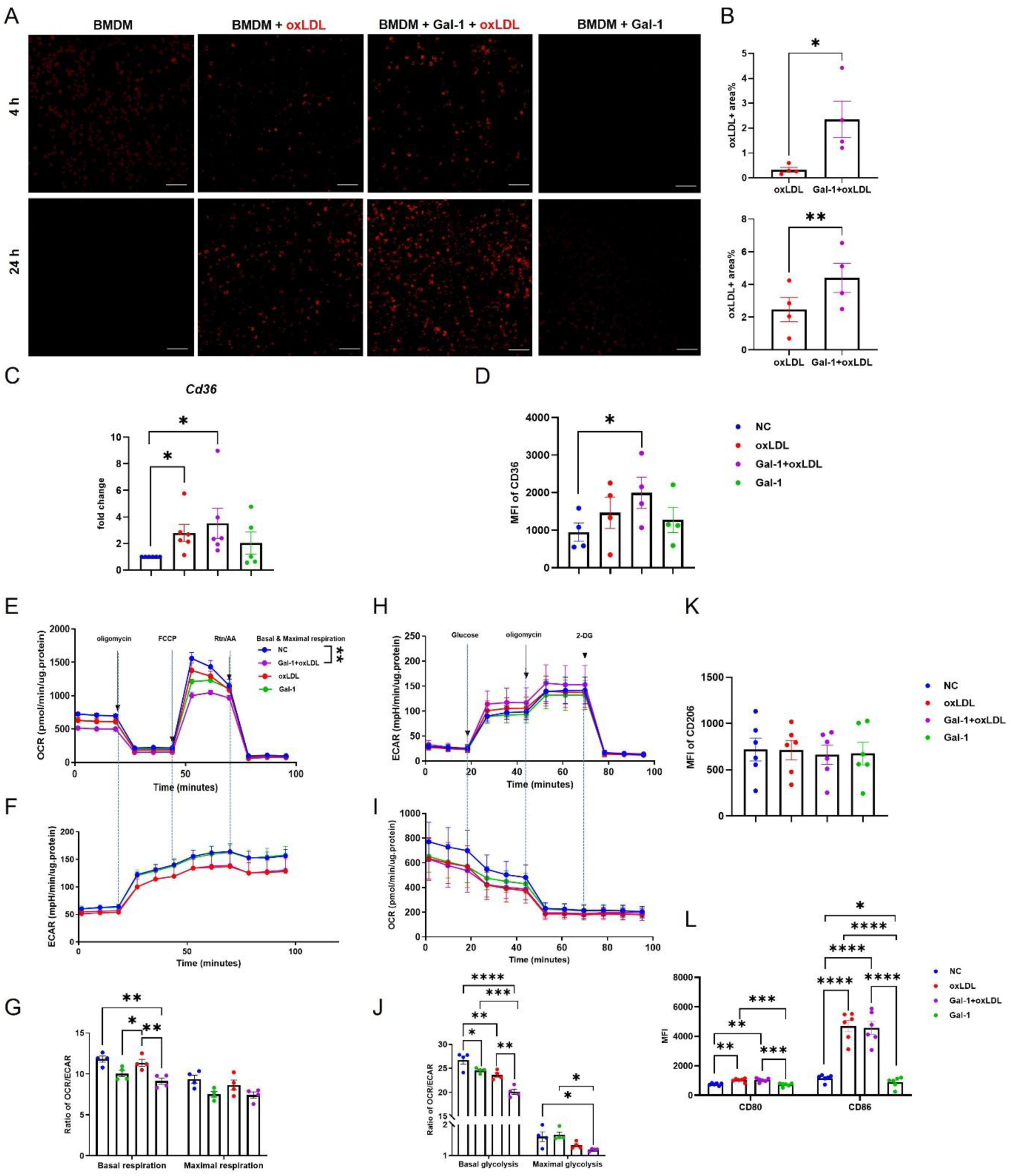
Effects of Gal-1 and oxLDL on energy metabolism and BMDM polarization. **(A-B)** BMDMs from *Apoe*^−/−^ mice were incubated for two distinct time periods (4 and 24 hours) with unlabelled oxLDL (50 µg/mL), with or without Gal-1 (100 nM). The formation of lipid droplets was then visualized through LipidSpot^TM^ 610 staining. **(A)** The images were captured using a confocal microscope (20 × magnification), and **(B)** lipid droplets were quantified as percent positive area of all cells. **(C)** The expression of *Cd36* in BMDMs was analysed by quantitative RT-PCR (n=5-6) and **(D)** surface expression by flow cytometry. **(E-J)** Oxygen consumption rate (OCR) and extracellular acidification rate (ECAR) of BMDMs were measured by the Agilent Seahorse XF Flux analyzer. Mitochondrial respiration **(E)** OCR, **(F)** ECAR and **(G)** OCR/ECAR ratio measured at basal conditions and after addition of FCCP to induce maximal respiration. Glycolysis **(H)** ECAR, **(I)** OCR, and **(J)** their ratio measured at basal conditions and after addition of oligomycin (ATP synthase inhibitor) to drive maximal glycolysis. Flow cytometry of **(K)** CD206, **(L)** CD80 and CD86 in BMDMs treated as indicated for 24 hours (n=6). Data are presented as the mean ± SEM. \**p* < 0.05, \*\**p* < 0.01, *\*\*\*p* < 0.001 and *\*\*\*\*p* < 0.0001. Statistical analysis: **(B-C)** two-tailed Student’s t-test against NC (negative control), **(D-L)** one-way ANOVA followed by Holm-Šídák’s post hoc test. Scale bar: 100 μm.

Recent studies indicate that cellular energy metabolism reprogramming is crucial for macrophage activation, influencing their phenotype and function. Glycolysis and mitochondrial respiration cannot be simply defined as pro– or anti-inflammatory [25]. To assess the impact of Gal-1 on energy metabolism, we measured oxygen consumption rate (OCR) and extracellular acidification rate (ECAR) in BMDMs treated for 24 hours with Gal-1 and/or oxLDL (**Fig.3E-J**). OxLDL reduced ATP-linked basal respiration (**Fig.3E**), consistent with previous reports, and Gal-1 also decreased basal respiration to a similar extent [26]. The combination of Gal-1 and oxLDL had an additive effect, suggesting independent mechanisms. The maximal respiration rate, stimulated by FCCP, followed a similar pattern to basal respiration, suggesting that the effects are likely due to impaired substrate supply or oxidation rather than ATP synthesis (**Fig.3E**) [27].

ECAR increased in response to electron transport chain inhibitors, a known compensatory response to mitochondrial oxidative phosphorylation (OXPHOS) inhibition (**Fig.3F**), but oxLDL independent of Gal-1 attenuated this increase [28]. The OCR/ECAR ratio was reduced by Gal-1 but not by oxLDL, indicating that Gal-1 shifts the metabolic balance toward glycolysis (**Fig.3G**). These findings were corroborated by the glycolysis stress test, where both Gal-1 and oxLDL reduced respiration, while inhibition of TCA/ATPase and glycolysis/hexokinase minimized OCR across all groups (**Fig.3H-J**). The OCR/ECAR ratio in the glycolysis stress test mirrored the mitostress test, further indicating a shift towards glycolysis in response to both treatments.

Further assessment of polarization markers showed that oxLDL upregulated surface expression of proinflammatory CD86 and CD80, and Gal-1 downregulated CD86 expression in untreated BMDMs. However, Gal-1 did not affect oxLDL-induced CD86 and CD80, suggesting that Gal-1 does not influence oxLDL-induced polarization (**Fig.3K, I**). Thus, atheroprotective effects of Gal-1 may occur independently of macrophage activation by oxLDL, or BMDMs may not be the ideal model for investigating these effects.

These data together indicate that Gal-1 enhances oxLDL uptake in BMDMs, potentially contributing to plaque foam cell formation and inflammation. However, Gal-1 failed to reverse the inflammatory profile induced by oxLDL, suggesting that its anti-atherogenic effects may be mediated through distinct inflammatory pathways. In addition, oxLDL-treated BMDMs as an entity in vitro may not fully recapitulate the phenotype of lesional macrophages in vivo.

### 3.5 Gal-1 promotes an anti-inflammatory phenotype in peritoneal foamy macrophages

To test the response to Gal-1 in macrophages other than BMDM, we focused on lipid-laden inflammatory macrophages, which more closely resemble the cellular environment in atherosclerotic lesions compared to BMDM. These macrophages were isolated from the peritoneal cavity following intraperitoneal injection of thioglycolate, in the context of WD-induced hypercholesterolemia (**Fig.4A**) [29]. In addition, we utilized a Gal-1 variant, Gal-1^E71Q^, which exhibits a near-complete loss of its ability to bind glycans, thus preventing cell aggregation (**Fig.S4A-D**) [30]. Mice were initially fed a WD for 2 weeks and subsequently treated with Gal-1, Gal-1^E71Q^ or PBS for an additional 2 weeks. After the entire 4-week treatment period, there were no significant differences in bodyweight gain and triglyceride levels (**Fig.S5A, B**). Plasma cholesterol levels remained unchanged (**Fig.S5C**), implying that 2 weeks of treatment was too short to affect cholesterol levels.

**Fig. 4.**
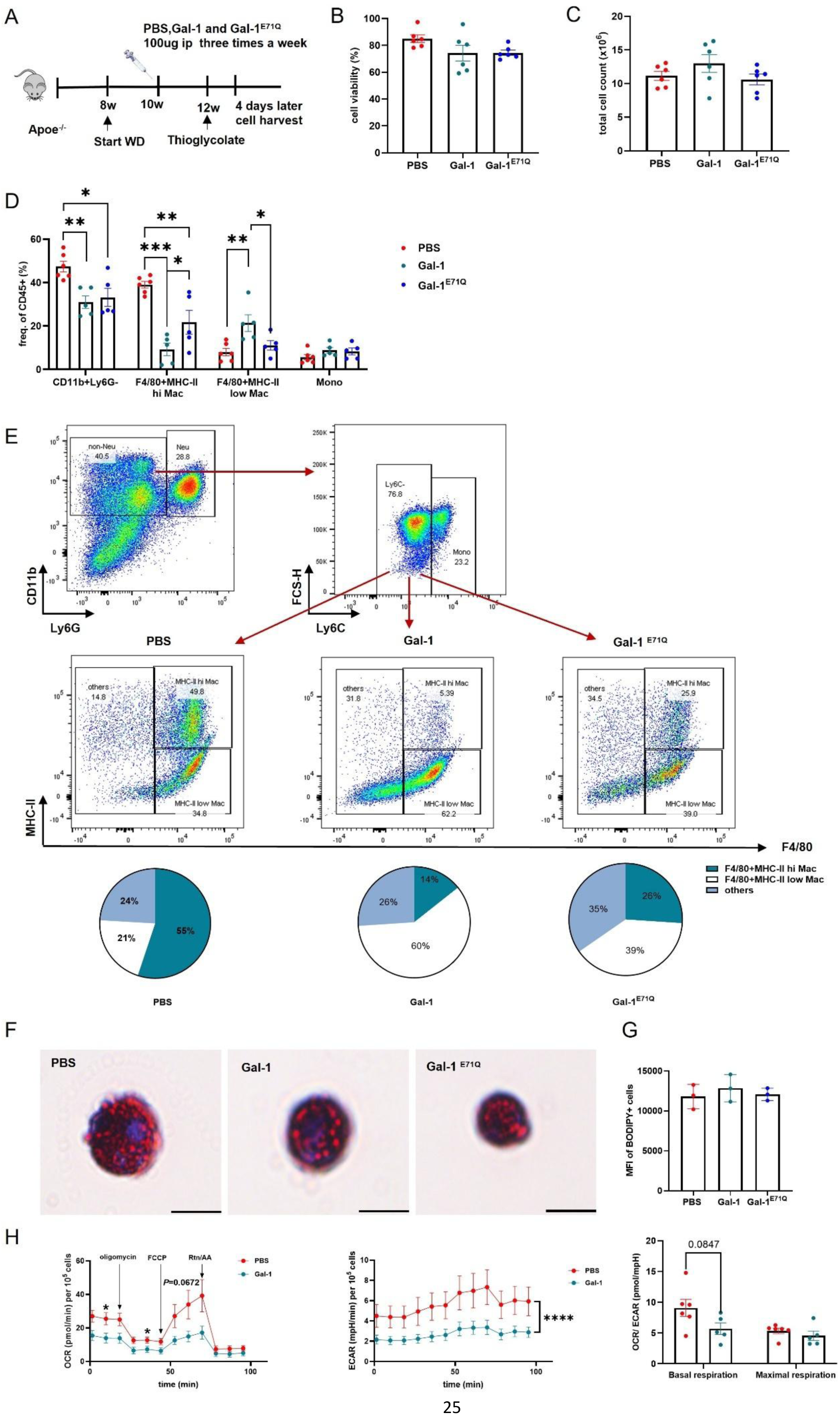
Effects of Gal-1 on peritoneal foam cells. **(A)** Experimental scheme of peritoneal foam cell induction. **(B)** Cell viability of peritoneal cells from PBS-, Gal-1-, and Gal-1^E71Q^-treated female mice (n=6 per group) was measured by Trypan blue staining. **(C)** Total number of cells in the peritoneal lavages per mouse was determined using an automated cell counter. **(D)** Frequency of myeloid cells were assessed by flow cytometry. **(E)** Flow cytometry gating strategy and corresponding pie chart depicting the relative abundance of CD11b⁺Ly6G⁻Ly6C⁻ cell subpopulations. **(F)** Representative images of peritoneal foamy macrophages, stained with Oil Red O (ORO). **(G)** Quantification of lipid droplets in peritoneal macrophages was obtained by flow cytometry after BODIPY staining (n=3). **(H)** Extracellular flux analysis of peritoneal macrophages from PBS– and Gal-1-treated mice after 4 weeks of WD (n=6). Mitochondrial respiration OCR, ECAR and OCR/ECAR ratio measured at basal conditions and after addition of FCCP to induce maximal respiration. Data are presented as the mean ± SEM. \**p* < 0.05, \*\**p* < 0.01 and *\*\*\*p* < 0.001. **(A-G)** one-way ANOVA followed by Holm-Šídák’s post hoc test, **(H)** each phase was analyzed using two-tailed Student’s t-test. Scale bar: 50 μm.

In the peritoneum, Gal-1 treatment did not affect cell viability (**Fig.4B**), and total peritoneal cell counts were similar across groups (**Fig.4C**). However, significant differences in cell composition were observed (**Fig.4D**). In controls, approximately half of the peritoneal cells were myeloid non-neutrophil cells (CD11b^+^Ly6G^-^) that predominantly consisted of inflammatory macrophages (F4/80^+^MHC-II^hi^), with a low number of homeostatic macrophages (F4/80^+^MHC-II^low^) and few monocytes. Treatment with Gal-1 and Gal-1^E71Q^ reduced the relative number of inflammatory macrophages, while increasing that of homeostatic macrophages (**Fig.4D, E**). The Gal-1^E71Q^ mutant induced similar effects, although the changes were less pronounced compared to Gal-1, suggesting that glycan-binding plays only a minor role and is not the sole cause of the observed effects on the macrophages. Furthermore, the frequency of B cells was reduced (**Fig.S5D**), while those of T cells and neutrophils were increased (**Fig.S5D, E**). The increase in neutrophils is in line with a previous report showing that Gal-1 mediates neutrophil influx during late stages of peritoneal inflammation [31]. These effects were largely dependent on carbohydrate recognition as the Gal-1^E71Q^ mutant behaved similarly to PBS control. There were no differences in the numbers of circulating leukocyte subsets between the groups (**Fig.S5F, G**). Although Gal-1 is known to induce (T cell) apoptosis [32], cell death cannot explain the observed effects given the comparable cell viability across groups, suggesting that Gal-1 may primarily influence macrophage proliferation and chemotactic factors. Collectively, these results indicate that Gal-1 treatment shifts the inflammatory profile of macrophages in an inflammatory environment, supporting an anti-inflammatory effect mediated by both sugar-binding dependent and independent mechanisms.

By accumulating lipids, macrophages exhibited foam cell-like characteristics (**Fig.4F**). However, lipid staining and flow cytometric quantification revealed no significant differences in intracellular lipid content between groups (**Fig.4G**), suggesting that Gal-1 does not influence peritoneal foam cell formation but rather reduces their inflammatory profile. This contrasts with findings in BMDMs, which may be attributed to differences in gene expression profiles between BMDMs and peritoneal macrophages in relation to foam cell formation [29].

Additionally, extracellular flux analysis showed that peritoneal macrophages from Gal-1-treated mice displayed a notable reduction in both OCR and ECAR but did not affect the OCR/ECAR ratio, suggesting a more quiescent metabolic phenotype, which might contribute to resolve inflammation (**Fig.4H**).

### 3.6 Transcriptomic profiling and polarization effect of Gal-1 on macrophages

To gain a deeper insight into the transcriptomic changes regulated by Gal-1 in macrophages, we isolated peritoneal inflammatory macrophages (F4/80^+^MHC-II^hi^) from mice as described in Figure 4A and conducted transcriptomic analysis by prime-seq [20]. We first investigated the effect of Gal-1 by analyzing differentially expressed genes (DEGs) between PBS– and Gal-1-treated macrophages. Out of 21,797 detected genes, we identified 450 DEGs, with 394 upregulated and 56 downregulated genes in the Gal-1-treated compared to the PBS-treated group (**Supplementary Table S1**). Among them, Gal-1 treatment upregulated the expression of genes associated with anti-inflammatory response (e.g., *Nlrp12, Rcor1, Trpc4*), engulfment functionality (e.g., *Trpc4, Marco*), and metabolism (e.g., *Suclg2, Dhdh, Cpt1b, Cpt2*) while downregulating apoptosis-related genes (e.g., *Siah2, Birc6, Wdr73*) (**Fig.5A**). Gene ontology (GO) analysis revealed enrichment in some inflammatory-related terms or pathways related to metabolism, such as Positive Regulation Of Transcription Of Notch Receptor Target (GO:0007221, *P*=0.0037), SREBP Signaling Pathway (GO:0032933, *P*=0.0076), Regulation Of Interleukin-18 Production (GO:0032661, *P*=0.01), Regulation Of Phagocytosis (GO:0050765, *P*=0.0343), Energy Reserve Metabolic Process (GO:0006112, *P*=0.0481), or Negative Regulation Of I-kappaB kinase/NF-κB Signaling (GO:0043124, *P*=0.0495).

**Fig. 5.**
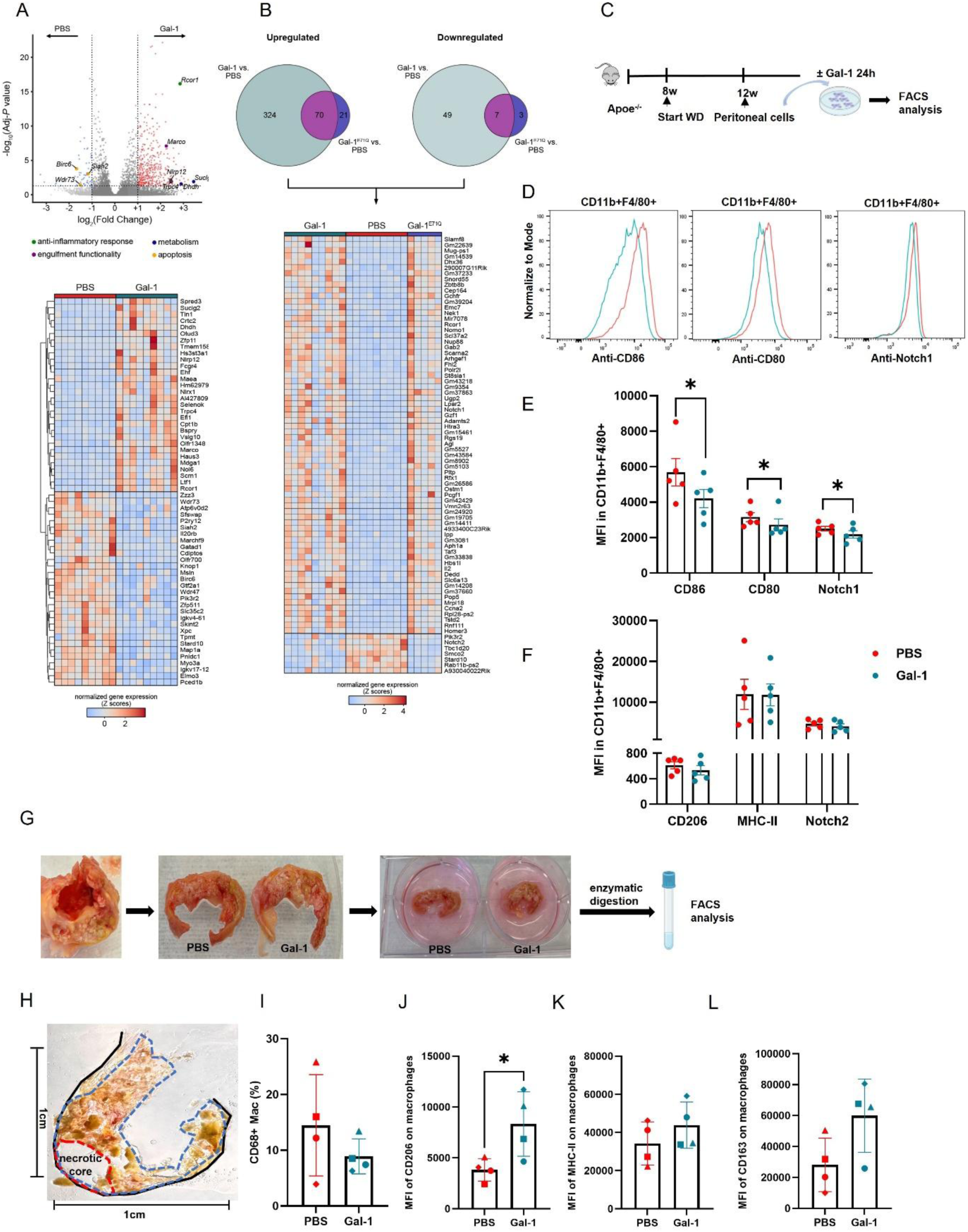
Transcriptional profiling and polarization effect of Gal-1 on macrophages. **(A)** Volcano plot (upper) displaying the differentially expressed genes (DEGs, adjusted *p* value < 0.05 and |log_2_FC| >1) of peritoneal macrophages from Gal-1-treated mice (n=9) against PBS-treated mice (n=9) as in Figure 4. Each colored dot represents an upregulated (red) or a downregulated (blue) DEG. Genes relevant to Gal-1 biological processes are highlighted and labeled. The heatmap represents the top 30 upregulated/downregulated genes. The complete list of DEGs is provided in ***Supplementary Table S1***. **(B)** Venn diagrams and the heatmap show the commonly upregulated and downregulated DEGs by both Gal-1 (n=9) and Gal-1^E71Q^ (n=5) vs. PBS (n=9). The complete list of DEGs between Gal-1^E71Q^ and PBS is provided in ***Supplementary Table S2***, the complete list of commonly regulated DEGs in Gal-1 and Gal-1^E71Q^ is found in ***Supplementary Table S3***. **(C-F)** Polarization effect of Gal-1 on peritoneal macrophages was assessed by flow cytometry. **(C)** Scheme of experimental design. **(D-F)** Surface expression of polarization markers (CD86, CD80, MHC-II, CD206) and Notch receptors (Notch1 and Notch 2) on peritoneal macrophages (CD11b^+^F4/80^+^) treated with 100 nM Gal-1 or PBS for 24 hours (n=5). **(G-L)** Effect of Gal-1 on human plaque macrophages. **(G)** Experimental work-flow: Atherosclerotic plaques from carotid/femoral endarterectomies were sliced into 2 mm thick sections and cultured for 24 hours with or without 100 nM Gal-1 treatment. Samples were then digested and analyzed by flow cytometry. **(H)** Representative image of human plaque section before culture. **(I)** The percentage of CD68^+^ macrophages from CD45^+^ cells. **(J-L)** Surface expression of CD206 (**J**), MHC-II **(K)** and CD163 **(L)** on macrophages was compared between groups. **(I–L)** Individual patients are represented by distinct symbols **(●, ▪, ▴, ◆)**. Data are presented as the mean ± SEM. \**p* < 0.05, two-tailed paired t-test on matched samples.

To dissect the role of glycan binding, we compared the DEGs in macrophages of mice treated with Gal-1^E71Q^, unable to bind glycans. Fewer DEGs (91 upregulated, 10 downregulated) were detected upon Gal-1^E71Q^ compared to PBS treatment (**Supplementary Table S2**), with 70 genes commonly upregulated and 7 genes commonly downregulated by both Gal-1 and Gal-1^E71Q^ (**Fig.5B and Supplementary Table S3**). These included genes relevant to regulating inflammatory response (e.g., *Rcor1, Slamf8, Pik3r2*) and the Notch signaling pathway (e.g., *Notch1, Notch2*), which has been implicated in the regulation of macrophage activation and polarization (**Fig.5B**) [33]. This observation may help to explain the shifted pattern of peritoneal inflammatory macrophages observed in Gal-1– and Gal-1^E71Q^-treated mice.

To further investigate the impact of Gal-1 on macrophage polarization, peritoneal cells were collected from mice after 4 weeks of WD (**Fig.5C**) and cultured with Gal-1 or PBS for 24 hours. Flow cytometry analysis revealed that Gal-1 treatment significantly reduced the expression of CD86 and CD80 on peritoneal macrophages. In addition, Gal-1 significantly downregulated Notch1 expression on macrophages and had no effect on CD206, MHC-II and Notch2 expression, suggesting an inhibition of pro-inflammatory macrophage phenotype (**Fig.5D-F**). Given that Notch1 is implicated in atherogenesis by regulating macrophage polarization and has been shown to interact with Gal-1, we speculate that Gal-1 may facilitate anti-inflammatory macrophage polarization through Notch1 suppression, thereby contributing to its atheroprotective effects [33, 34].

To further validate the polarization effect of Gal-1 on macrophages within atherosclerotic plaques, we cultured freshly isolated plaques obtained from femoral or carotid artery endarterectomy samples [35]. Fresh plaque was sliced into 2mm thick sections and placed on a wetted sponge at the air-medium interface, followed by treatment with or without 100 nM Gal-1 for 24 hours (**Fig.5G**). To visualize the baseline plaque structure, one slice was further cryosectioned into 10 µm sections, and images were captured prior to treatment (**Fig.5H**). After 24 hours, the plaques were enzymatically digested and cell populations were analyzed by flow cytometry. While there was no noticeable change in the proportion of total macrophages (**Fig.5I**), the expression of CD206 in macrophages was significantly upregulated following Gal-1 stimulation (**Fig.5J**). However, the expression of MHC-II and CD163 remained unaltered (**Fig.5K, L**). These observations further illustrate the anti-inflammatory action of Gal-1 in modulating macrophage inflammatory reprogramming within atherosclerotic plaques.

### 3.7 Gal-1 skews T helper balance and boosts Th17-Treg immunity

Gal-1 has been previously reported to modulate T-cell mediated immune responses [18]. To assess the impact of Gal-1 injection on T cell profiling in our atherosclerosis model, we conducted plasma cytokine analysis from mice, as shown in Figure 1. However, no significant differences were observed for Th1-(IFN-γ, TNF-α) or Th2-related cytokines (IL-4, IL-5, IL-9, and IL-13) between PBS– and Gal-1-treated mice. Intriguingly, several cytokines associated with the Th17 immune response (IL-22, IL-23, IL-17, IL-6), which are typically linked to pro-inflammatory responses, were elevated in Gal-1-treated mice (**Fig.6A**). This finding suggests that Gal-1 may exert complex immunomodulatory effects, potentially enhancing Th17 signaling pathways. Notably, concurrent IL-17 and IL-10 elevation may indicate a regulatory Th17 response associated with plaque reduction [36].

**Fig. 6.**
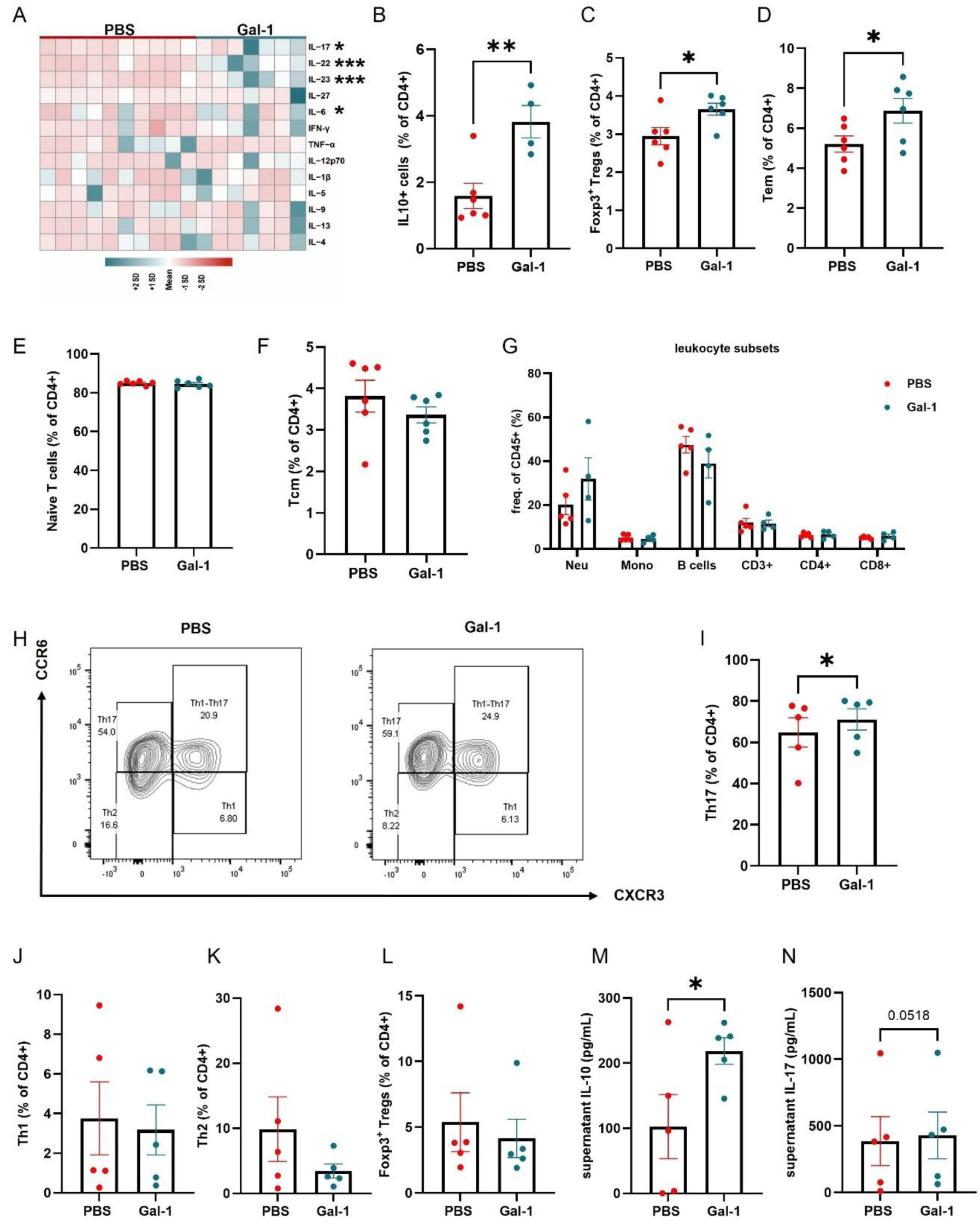
Plasma cytokine profiling and immune cell analysis. **(A)** Heatmap representing plasma multiplex cytokine analysis from PBS-(n=10) and Gal-1-treated *Apoe*^−/−^ mice (n=7). **(B-F)** Flow cytometry was used to quantify the proportions of distinct T cell subsets within the CD4⁺ T cells in blood. (**B**) IL-10-secreting CD4⁺ T cells, **(C)** regulatory T cells (Treg: CD4⁺Foxp3⁺), (**D**) effector memory T (Tem: CD44⁺CD62L⁻), (**E**) naïve T (CD44⁻CD62L⁺), (**F**) and central memory T cells (Tcm: CD44⁺CD62L⁺) cells (n=4-6 per group). **(G)** Proportions of distinct leukocyte subsets within CD45^+^ cells in blood. **(H-N)** CD4^+^ T cells were isolated by negative selection from spleens and cultured for 72 hours with 50 nM Gal-1 or PBS. **(H)** Gating strategy for T helper subsets, **(I)** percentage of Th17 (CCR6^+^CXCR3^-^), **(J)** Th1 (CCR6^-^CXCR3^+^), **(K)** Th2 (CCR6^-^CXCR3^-^), **(L)** Treg (CD4^+^Foxp3^+^). **(M)** IL-10 and **(N)** IL-17 levels in the supernatant were measured using Elisa (n=5 per group). **(A)** Mann-Whitney U-test. Data in **(B-N)** are shown as mean ± SEM. \**p* < 0.05, \*\**p* < 0.01 and *\*\*\*p* < 0.001, (**B-G**) two-tailed Student’s t-test. (**I-N**) two-tailed paired Student’s t-test on matched samples.

To investigate the cellular source of IL-10 production, we analyzed leukocyte and T-cell subsets in the blood. Gal-1 treatment significantly increased the percentage of IL-10–producing CD4^+^ T cells in the blood (**Fig.6B**), while no changes were observed in inguinal lymph nodes (**Fig.S6A**). The increase in IL-10–producing CD4^+^ T cells was accompanied by an expansion of CD4^+^FoxP3^+^ regulatory T cells (Tregs) (**Fig.6C**) and a significant increase in effector memory CD4^+^ T cells (Tem, CD44^+^CD62L^-^) (**Fig.6D**) in the blood of Gal-1-treated mice. As both Tregs and effector memory CD4⁺ T cells can produce IL-10, their expansion likely contributes to the elevated frequency of IL-10–producing CD4⁺ T cells observed in Gal-1–treated mice [37]. However, the frequencies of naïve (CD44^-^CD62L^+^) and central memory (Tcm, CD44⁺CD62L⁺) CD4⁺ T cells, as well as other leukocyte subsets, remained unchanged (**Fig.6E-G**). Leukocyte composition and CD4⁺ T-cell apoptosis in the spleen and lymph nodes were also comparable between groups **(Fig.S6A, B**).

To further elucidate the effects of Gal-1 on T cell differentiation, CD4^+^ T cells were isolated from the spleens of *Apoe*^−/−^ mice and cultured in vitro with or without Gal-1 for 72 hours. After three days of culture, Gal-1 treatment significantly skewed T cell differentiation toward the Th17 subset, while the frequencies of Th2, Treg, and Th1 population remained unchanged (**Fig.6H-L**), indicating that Gal-1 preferentially promotes the differentiation of naïve T cells into Th17 cells. Analysis of cytokines in the supernatant revealed a marked increase in IL-10 production, while IL-17 levels remained unchanged in Gal-1-treated cells (**Fig.6M, N**). These findings suggest that Gal-1 modulates the equilibrium between Treg and Th17 responses, potentially favoring a regulatory Th17 phenotype that may contribute to the attenuation of atherosclerosis.

## 4. Discussion

This study offers new insights into the role of Gal-1 in atherosclerosis, revealing its capacity to reduce atherosclerotic burden and promote less inflammatory plaque formation. These effects were associated with changes in macrophage immunometabolism and T cell immunity after Gal-1 treatment (**Fig.7**). In macrophages, Gal-1 induced a more quiescent metabolic state, and reduced inflammatory activity. The use of the glycan-binding-deficient variant Gal-1^E71Q^ demonstrated that the anti-inflammatory effects on macrophages are mediated only partially through its carbohydrate-binding capacity. Furthermore, Gal-1 induced a distinct cytokine profile in blood, characterized by elevated levels of Th17 immune response associated cytokines. This was accompanied by an expansion of Tregs and a shift in T-cell differentiation towards the Th17 phenotype.

**Fig. 7.**
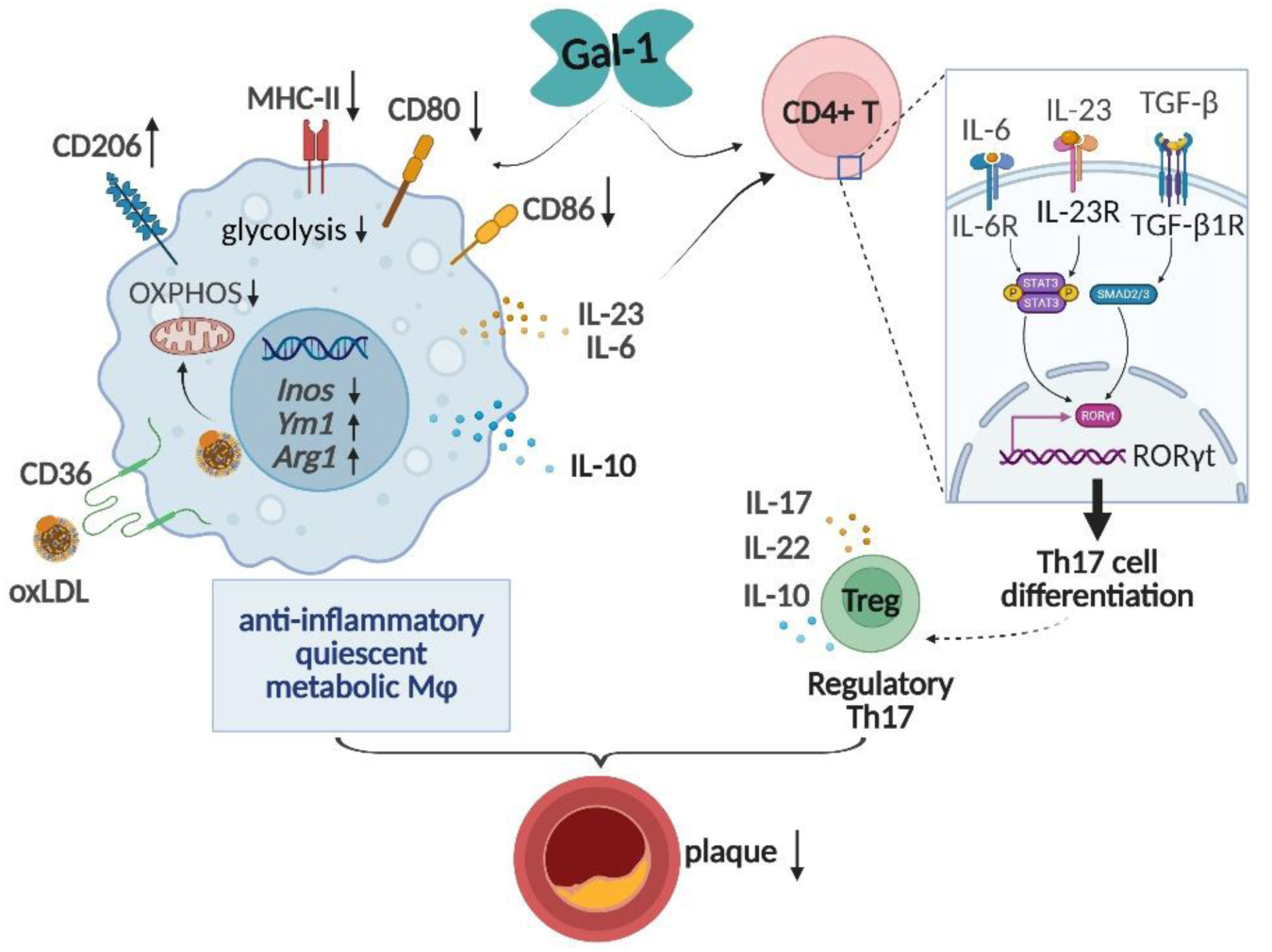
Gal-1 reprograms macrophage metabolism and shapes T cell responses to reduce atherosclerosis. Gal-1 acts on macrophages to induce an anti-inflammatory, metabolically quiescent phenotype characterized by increased CD206, reduced MHC-II, CD80, and CD86 expression, decreased glycolysis and OXPHOS, and altered gene expression (↓*Inos*, ↑*Ym1*, ↑*Arg1*). Gal-1 also modulates CD4⁺ T cell responses by shaping cytokine signals that promote a regulatory Th17 profile with increased IL-17, IL-22, and IL-10 supporting a regulatory phenotype. The combined effects of macrophage reprogramming and T cell modulation likely contribute to reduced plaque development.

These findings further support and build upon previous research, demonstrating the atheroprotective effects of Gal-1 associated with smooth muscle cell functions, whereas the atherogenic impact on macrophages and T cells remains less clear [16].

Our study showed that Gal-1 treatment elevated plasma IL-10 levels and increased IL-10-secreting T cells, correlating with reduced plaque size. Gal-1 has previously been shown to enhance IL-10-mediated T cell tolerance and to suppress autoimmune neuroinflammation, with IL-10 widely recognized as a potent anti-inflammatory cytokine [38]. Yet the role of IL-10 in atherosclerosis remains controversial: deleting the IL-10 receptor in myeloid cells reduces lesion size and IL-10 boosts oxLDL uptake and foam cell formation [39, 40]. We hypothesized that Gal-1 could reduce atherosclerosis in an IL-10-dependent manner, potentially explaining the reduced cholesterol levels we observed. However, neutralizing IL-10 with an established monoclonal antibody led to a reduction in lesions, and when combined with Gal-1, it failed to restore the reduced atherosclerotic burden. Instead, we observed an additive and thus mechanistically likely independent reduction. One explanation could be that the antibody treatment effects differ from the results of genetic models, as some compartments (e.g., intracellular) might not be antibody accessible. Therefore, if the functions of IL-10 are context-dependent, blocking only a portion of its activity may have led to the opposite effect seen with genetic deletion. Another key factor may be the cholesterol levels, which were reduced by both IL-10 Ab and Gal-1 but showed no additive effects when combined, suggesting a dependency between these two effects. Nevertheless, despite a reduction in plaque size, the neutralization of IL-10 promoted features of plaque instability that are more in line with the anticipated anti-inflammatory effects of IL-10.

Gal-1-treated mice exhibited lower plasma cholesterol levels after 4 weeks, though the underlying mechanisms are complex. Gal-1-deficient mice have been reported to downregulate PPARγ-dependent genes involved in gluconeogenesis and lipogenesis, with reduced liver triglyceride accumulation [41]. However, Gal-1-treated mice did not affect cholesterol accumulation in the liver nor change the key lipid metabolism genes, suggesting a liver-independent effects on cholesterol. Intestinal cholesterol absorption was not a factor either. Moreover, Gal-1 is known to bind extracellular vesicles [24], and hypothetically also lipoprotein particles but it did not precipitate lipoprotein particles in our study. Another possibility is enhanced oxLDL uptake by macrophages as observed in BMDMs, which might have partially contributed to reduced plasma cholesterol levels. Possibly, several small factors are the reason which make the identification of a dominant pathway difficult.

Macrophage metabolic reprogramming plays a pivotal role in driving the inflammatory response, with evidence linking this process to oxLDL/CD36 signaling and is essential in the activation and polarization of macrophages, thereby influencing the progression of atherosclerosis [42]. Building on this, we further investigated the metabolic and inflammatory reprogramming of macrophages following Gal-1 treatment. Gal-1 enhanced oxLDL uptake in BMDMs possibly through upregulation of CD36. However, the effects of Gal-1 on energy metabolism were independent of oxLDL. Specifically, Gal-1 modestly reduced OXPHOS in BMDMs, accompanied by a shift toward glycolysis, and more profoundly reduced both respiration and glycolysis in peritoneal foam cells, indicating a quiescent phenotype as in glioma metabolism [43]. The effect of Gal-1 on the energy metabolism might be essential for anti-inflammatory macrophage activation. Indeed, polarization markers indicated an alternatively activated phenotype mediated by Gal-1 in BMDMs and peritoneal foamy macrophages. Additionally, Gal-1 injection converted the high percentage of pro-inflammatory to anti-inflammatory macrophages, suggesting that Gal-1 inhibits pro-inflammatory macrophage programming. This immunometabolic profile of Gal-1-treated macrophages, together with the increase in anti-inflammatory macrophages in the plaques of Gal-1-treated mice, is consistent with previous research showing that Gal-1 supports a switch to a pro-resolving macrophage phenotype under inflammatory conditions [11, 44]. The glycan-binding defect mutant Gal-1^E71Q^ yielded similar anti-inflammatory effects suggesting a partially glycan-independent mechanism [30]. Gal-3 can modulate inflammation via interaction with chemokines in a carbohydrate-binding independent manner [9]. A similar mechanism may apply to Gal-1, explaining the observed effects of Gal-1^E71Q^. Interactions with chemokines may be one of many possibilities of how galectins can influence inflammation in a carbohydrate-binding independent manner.

The transcriptomic profile of inflammatory peritoneal foam cells from Gal-1 or Gal-1^E71Q^-treated mice aligns with this. The number of differentially expressed genes was much higher in the Gal-1 group than in the Gal-1^E71Q^ group and comprised most of the genes regulated by Gal-1^E71Q^. It remains speculative which of the pathways are responsible for the anti-inflammatory and immunometabolic effects on macrophages and atherosclerosis. We observed that some pathways, such as the Notch-signaling pathway, are commonly affected by both variants and are known to play an inflammatory role, further indicating the anti-inflammatory effects of Gal-1 through glycan-independent mechanisms.

On the other hand, T cells are strongly involved in atherogenesis, and Gal-1 is implicated in several T cell responses such as suppressing Th1 responses, promoting a Th2-skewed immune profile, and inducing T cell apoptosis [18, 32]. However, our study did not observe significant alterations in Th2/Th1-associated cytokines or T cell apoptosis. Instead, we noted an increase in IL-10 production and a higher frequency of IL-10-secreting CD4^+^ T cells along with an expanded pool of effector memory T cells and Tregs, indicating that Gal-1 might modulate Treg function, which is consistent with studies showing that Gal-1-deficient mice exhibited reduced regulatory activity in CD4^+^CD25^+^ T cells that are endowed with atheroprotective functions [45, 46]. The observed cytokine profile indicates that IL-17 release likely results from macrophage activation, which triggers IL-23 and IL-6 production, promoting Th17 differentiation and elevating IL-17 and IL-22 levels [47]. Consistent with this, Gal-1 enhanced the differentiation of in vitro cultured T cells into Th17 cells. These findings suggest that Gal-1 may modulate T cell responses at concentrations below its apoptotic threshold, allowing it to drive specific subsets without inducing cell death. Indeed, it has been reported that Gal-1 does not bind to naïve T cells, Th2 cells, or resting Tregs but strongly binds to Th17, and activated Tregs, where it induces apoptosis only at micromolar concentrations [48].

Direct effects of Gal-1 on T cell differentiation into Th17 cells that are carbohydrate dependent can be mediated by CD69 have been reported [49].

Our results indicate that Gal-1 enhances a Th17 response. While IL-17 is generally regarded as a pro-inflammatory cytokine, its functional role—whether pro-inflammatory or homeostatic—appears context-dependent [50]. Notably, IL-17 has also been shown to stabilize atherosclerotic plaques and exhibit anti-inflammatory properties when combined with elevated IL-10 levels, as observed in our study [36]. The subset of T helper cells that produce both IL-10 and IL-17 has been described as regulatory Th17 cells, highlighting the balance between homeostatic and pathogenic IL-17-producing cells, as previously reviewed [36, 51]. Furthermore, Th17 cells have been reported to transdifferentiate into Tregs during the resolution of inflammation [52]. It is therefore possible that we observed an intermediate stage in which IL-17-producing T cells are transdifferentiating into IL-10-producing T cells under the influence of Gal-1.

Several limitations of this study should be acknowledged. First, although Gal-1 treatment was associated with a reduction in plasma cholesterol, the precise mechanisms driving this systemic lipid-lowering effect remain unclear. However, the modest decrease in cholesterol levels suggests only a limited impact on atherosclerotic lesion development. In addition, while there was no difference in sex-based atherosclerotic plaque reduction after Gal-1 treatment in this study, the small sample size may have limited the statistical power to detect potential sex-related effects. Moreover, while our data indicate that Gal-1 exerts immunomodulatory effects that are at least partly independent of IL-10, and may influence T cell subsets such as Th17 and Treg populations, these observations require further validation. Finally, our investigation focused on immune cells. Other cell types such as smooth muscle cells may also contribute but were not investigated here. Despite these limitations, our findings demonstrate a link between Gal-1 treatment and immunomodulatory effects in the context of atherosclerosis, supporting its potential as an immunoregulatory therapy for this disease.

## CRediT authorship contribution statement

**Ya Li and Julian Leberzammer:** Investigation, Methodology, Writing-original draft. **Xavier Blanchet, Rundan Duan, Rui Su, Michael Lacy, Veit Eckardt, Yvonne Jansen and Yvonne Döring:** Investigation, Methodology. **Vasiliki Triantafyllidou and Donato Santovito:** Software, Formal analysis, Visualization, Data Curation. **Michael Hristov:** Resources, Methodology. **Eva Briem and Wolfgang Enard:** Resources, Software, Project administration. **Anna Simone Jung, Joel Guerra and Alexander Bartelt:** Resources, Investigation. **Jürgen Bernhagen, Christian Weber and Dorothee Atzler:** Resources, Funding acquisition. **Herbert Kaltner and Anna-Kristin Ludwig:** Resources, Conceptualization. **Philipp von Hundelshausen:** Conceptualization, Funding acquisition, Supervision, Writing-review & editing.

## Ethics approval

All animal procedures were performed conforming to the guidelines from Directive 2010/63/EU, die Regierung von Oberbayern, Germany; (ROB-55.2Vet-2532.Vet_02-17-180, ROB-55.2-2532.Vet_02-22-150), the ARRIVE guidelines and the ethical standards laid down in the 1964 Declaration of Helsinki and its later amendments.

## Acknowledgments

C.W. is a Van de Laar professor of atherosclerosis. The graphical abstract was created with BioRender.com.

## Supplementary Information

### Supplemental Methods

#### Expression and purification of Gal-1 and Gal-1^E71Q^

For recombinant protein expression the pGEMEX-1 expression plasmid and the pGEX-6p2 were used for Gal-1 and its mutant variants Gal-1^E71Q^, respectively. The pGEMEX-Gal-1 or the pGEX-Gal1^E71Q^ plasmid was transformed into Escherichia coli BL21 (DE3)-pLysS (Promega) for recombinant production. Cells were grown in LB (Roth, Karlsruhe, Germany) medium containing the appropriate antibiotic for 16 h at 37 °C and then transferred to TB (Roth) medium. After initial growth for 2-3 h at 37 °C up to an OD600nm of 0.6-0.8, gene expression was induced using 100 µM β-1-thio-D-galactopyranoside (IPTG), and bacteria were cultured at 37 °C (Gal-1) or 22°C (Gal-1^E71Q^) for additional 16 h. The protein was purified from bacterial extracts by affinity chromatography on lactosylated Sepharose 4B (Gal-1) or glutathione Sepharose (Gal-1^E71Q^, Cytiva, Dreieich, Germany) as described before [1]. After chromatographic purification the buffer was exchanged to 10 mM PBS pH 7.2 via PD10 columns (Cytiva, Dreieich, Germany). Purity was ascertained by one-dimensional gel electrophoresis under denaturing conditions.

### Multiplex immunoassays

The ProcartaPlex™ Mouse Cytokine Kit (Invitrogen™, EPX170-26087-901) was used to quantify cytokines in plasma according to the manufacturer’s protocol. Cytokine levels were measured using the Luminex™ xMAP platform, and data were analyzed with ProcartaPlex™ Analyst software (Thermo Fisher Scientific).

### In vitro CD4+ T cell differentiation

Untouched CD4^+^ T cells were isolated from the spleen of *Apoe*^−/−^ mice by depleting non-target cells using magnetic beads (Miltenyi Biotec) and seeded at 1 × 10^5^ cells/well. Cells were stimulated with Dynabeads™ Mouse T-Activator CD3/CD28 (Invitrogen) at a 1:1 bead-to-cell ratio, with or without 50 nM Gal-1, in RPMI-1640 medium supplemented with 10% FBS, 2 mM L-glutamine, 1% Penicillin/Streptomycin, 10 mM HEPES, 1% MEM non-essential amino acids, 1 mM sodium pyruvate, and 50 µM beta-mercaptoethanol (all Gibco). After three days, supernatants were collected for IL-17 and IL-10 measurement (DuoSet ELISA, R&D Systems), and cell pellets were analyzed by flow cytometry following bead removal.

### Histology and immunofluorescence

Hearts, including the aortic roots and aortic arches with the main branch points (brachiocephalic artery, left subclavian artery, and left common carotid artery), were dissected and fixed in 4% paraformaldehyde (PFA). After fixation, samples were embedded in paraffin and sectioned at 4 µm thickness. Slides were deparaffinized and stained using an Elastin/van Gieson (EVG) staining kit (Morphisto, 12739) to assess lesion size and necrotic core. Fibrous cap was defined as the eosinophilic layer overlaying the necrotic core of the plaque. Ten equally dispersed measurements of cap thickness were taken from the most vulnerable necrotic core of each plaque section, and the averages were calculated for each mouse. For the aortic roots, serial sections from the proximal to distal portions were used to evaluate plaque formation. Lesion size in the aortic arches was quantified after EVG staining of 4 µm transverse sections, and averages were calculated from 3-4 sections. Plaque collagen content was evaluated using Picro Sirius Red staining (Sigma-Aldrich, P6744-1GA), with averages calculated from 3 sections. Neutral lipid accumulation in the liver was assessed using Oil-Red-O (ORO) staining on cryosections. Liver tissue was embedded in Tissue-Tek O.C.T. compound (Sakura) for cryosectioning. ORO-positive areas were quantified in 6 µm sections, and averages were calculated from 3 sections.

For immunofluorescence staining, aortic root sections were dewaxed and rehydrated, followed by antigen retrieval and blocking with either 10% goat serum or 5% BSA. Sections were incubated overnight at 4 °C with primary antibodies against Mac-2, iNos, CD206, CD3, and perilipin-2. After washing, sections were incubated with secondary antibodies conjugated with 4’,6-diamidino-2-phenylindole (DAPI) for nuclear staining in the dark for 1 hour at room temperature (**antibodies listed in Major Resources table S1**). Slides were washed and mounted with DAKO fluorescence mounting medium (S3023, Agilent) for microscopy.

Digital images were captured using a Leica microscope (Leica, Germany). Image analysis was performed in a blinded manner using Leica Application Suite LAS V4.3 software and ImageJ software. For each marker, 3 to 5 cross-sections per mouse were quantified, and the average was calculated.

### Flow cytometry

Blood was collected from the posterior orbital vein into citrate tubes, and inguinal lymph nodes, bones, and spleen were harvested in PBS and kept on ice. The spleen and lymph nodes were mechanically mashed using a 50 µm cell strainer (Cell-Trics, Partec) with FACS buffer (2% FBS in PBS), and bone marrow cells were flushed out. The cell pellets from the spleen, bone marrow, peritoneal lavage, and blood were lysed with ACK buffer (0.15 mM NH_4_Cl, 1mM KHCO_3_, 0.1 mM Na_2_EDTA, pH 7.2-7.4) to eliminate red blood cells. The remaining cell suspensions were stained with live/dead fixable viability dyes (Invitrogen) for 10 minutes, followed by Fc blocker (anti-CD16/CD32, 1:200, Biolegend) for 10 minutes. After a wash with FACS buffer, the cells were stained with surface antibodies (**Major Resources table S2**) for 45 minutes in the dark at 4 °C. Peritoneal lavage samples were analyzed using a BD FACSAria III Cell Sorter (BD Biosciences) to isolate inflammatory peritoneal macrophages (CD45^+^CD11b^+^F4/80^+^MHC-II^+^). Sorted cells were stored in RLT buffer (Qiagen) with 1% beta-mercaptoethanol at –80 °C.

For intracellular staining, cells were permeabilized using the permeabilization buffer (eBioscience, 00-5521-00) according to the manufacturer’s instructions before antibody staining. Apoptosis was analyzed using the Apoptosis Detection Kit (Biolegend, 640934).

For intracellular cytokine staining (IL-10, INF-γ, IL-4, and IL-17), single-cell suspensions from blood, spleen, and lymph nodes were incubated with Cell Stimulation Cocktail (1:500 dilution, eBioscience) and Brefeldin A (1:1000 dilution, Biolegend) in 200 µL of T-cell culture medium (RPMI-1640 with 10% FBS, 2 mM L-glutamine, and 1% Penicillin/Streptomycin, all from Gibco by Life Technologies) for 5 hours at 37 °C. The cell suspensions were then washed with PBS, followed by Fc blocker incubation (anti-CD16/CD32, 1:200, Biolegend) for 10 minutes. Cells were stained with surface antibodies for 45 minutes in the dark at 4 °C. After a wash, live/dead staining was performed using Fixable Viability Dyes (Invitrogen) in the dark for 20 minutes at 4 °C. The cells were washed again, resuspended in FACS buffer, and analyzed by flow cytometry.

For intracellular lipid measurement in peritoneal macrophages, isolated cells were fixed with 2% PFA at room temperature for 20 minutes, washed with PBS, and stained with 10 µM BODIPY™ 493/503 (Invitrogen) at room temperature for 30 minutes. After washing, cells were analyzed by flow cytometry.

Flow cytometry measurements were performed using a BD FACS Canto II (BD Biosciences) and BD LSRFortessa (BD Biosciences). Data were analyzed with FlowJo v10.6.2 software (Tree Star, Inc). Absolute cell counts were determined using CountBright absolute counting beads (Invitrogen, C36950).

### Bone marrow-derived macrophage (BMDM) experiments

Bone marrow cells were flushed from the femur or tibia using cold PBS, and the total bone marrow progenitors were counted. The cells (1.5 x 10^5^ cells/mL) were then seeded into culture plates containing macrophage complete medium (DMEM/F12-10 medium, 10% FBS, 10% L929-conditioned medium, 50 µg/mL Gentamicin). On day 4, the medium was replaced with a 40% old medium and 60% fresh medium mixture, and the cells were cultured until day 7 to allow for differentiation into BMDMs.

For oxLDL uptake experiments, BMDMs (1 × 10^5^ cells/well) were seeded onto chamber slides (Ibidi, 80427) and incubated with 50 µg/mL oxLDL (Invitrogen™, L34357), with or without 100 nM Gal-1 for either 4 hours or 24 hours at 37 °C. After washing with cold PBS, intracellular oxLDL was stained with LipidSpot™ 610 (Biotium, 70069) for 20 minutes, fixed with 2% PFA for 15 minutes. Confocal microscopy was used to capture five random fields per sample, ensuring consistent cell density across samples. The average area of LipidSpot-positive staining was analyzed using ImageJ. Polarization of BMDMs was assessed by flow cytometry.

### Extracellular flux analysis

Oxygen consumption rate (OCR) and extracellular acidification rate (ECAR) were measured using the Seahorse Cell Mito Stress and Glycolysis Stress Tests on an XF-24 Extracellular Flux Analyzer (Agilent). Briefly, BMDMs (1 × 10^5^ cells/well) were seeded into 24-well seahorse cell culture plate after 7 days of differentiation. Subsequently, the cells were treated with 100 nM Gal-1 alone or in combination with 50 µg/mL ox-LDL for 24 hours. Prior to measurement, the cells were treated and washed according to the manufacturer’s protocol. For Mito Stress Test, three replicates of data were obtained under basal conditions and on addition of oligomycin (1.5 µM), fluoro carbonyl cyanide phenylhydrazone (FCCP, 1.5 µM), and rotenone + antimycin A (Rtn/AA, 2.5 µM) (Sigma-Aldrich). For glycolysis test, three replicates of data were obtained under basal conditions and on addition of glucose (10 mM), oligomycin (1.5 µM), and 2-Deoxy-D-glucose (2-DG, 50 µM). After the measurement, cells were lysed, and protein concentrations were determined using the BCA Protein Assay Kit (Thermo Scientific™ 23225). All metabolic rates were normalized to total protein levels.

For peritoneal foamy macrophage Mito Stress Test, peritoneal lavage was collected, and erythrocytes were removed by ACK buffer. B cells were then depleted using mouse CD19 MicroBeads Isolation Kit (Miltenyi Biotec), followed by neutrophil depletion with anti-mouse Ly6G Dynabeads™ (Thermo Fisher). Seahorse plates were pre-coated with Poly-D-Lysine (Sigma) (1:10 dilution) for 1 hour at RT, then rinsed and dried. After isolation, 1 × 10^5^ cells/well were seeded in 24-well Seahorse plates (Agilent) in RPMI-1640 (10% FBS, 2 mM L-glutamine, 1% Penicillin/Streptomycin) (all Gibco) and incubated for 4 hours at 37 °C prior to Seahorse measurements. Triplicates of data were taken under basal conditions and on addition of oligomycin (1.5 µM), FCCP (1.5 µM), and Rtn/AA (2.5 µM) (Sigma-Aldrich). After the readout, all metabolic rates were normalized to cell numbers.

### Gene expression analyses

Total RNA was extracted using RNeasy mini kit (Qiagen) and reverse transcribed (Applied Biosystems™, 4368814) according to the manufacturer’s instructions. Real-time quantitative PCR (qPCR) was performed on the QuantStudio™ 7 Pro Real-Time PCR System (Thermo Fisher) using the TaqMan™ Gene Expression Master Mix (Applied Biosystems™, 4369016). Primers and probes were purchased from Integrated DNA Technologies (IDT) and TaqMan **(Major Resources table S3)**. Target gene expression was normalized to 18S ribosomal RNA (Applied Biosystems™, 4332641) gene expression and represented as fold change relative to the control group.

### RNA sequencing and data analysis

Inflammatory peritoneal macrophages were sorted from mice as described above. RNA-sequencing was conducted using the prime-seq method developed by Janjic et al. (2022) [2]. The complete protocol for prime-seq, including primer sequences, is available at protocols.io (<u>prime-seq (protocols.io)</u>). Briefly, cDNA synthesis was performed using Maxima H Minus reverse transcriptase, oligo-dT primer E3V7NEXT, and template switching oligo. After pooling, remaining primers were removed with Exonuclease I. Subsequently, cDNAs were pre-amplified using KAPA HiFi HotStart polymerase. Libraries were prepared using the NEBNext Ultra II FS DNA Library Prep Kit for Illumina (New England Biolabs, E7805S) with a custom ligation adapter, and dual indexing with TruSeq i5 and Nextera i7 index primers. Library concentrations were quantified before sequencing utilizing an Agilent 2100 Bioanalyzer instrument. Sequencing was performed on an Illumina NextSeq 1000/2000 instrument, with read lengths of 28 bases (read 1), 8 bases (read 2), 8 bases (read 3), and 93 bases (read 4). Quality assessment of the fastq data files was conducted using fastqc (v0.11.5), followed by polyA trimming using cutadapt (v 4.1). Subsequent read filtering, mapping, and counting were performed using the zUMIs pipeline (v 2.9.7), with reads mapped to the mouse genome (mm10). Differential gene expression analysis (DEA) was calculated using DESeq2 (v. 1.44.0), a Bioconductor package in R (v. 4.4.2). Differentially expressed genes (DEGs) were identified by criteria: adjusted two-tailed *P* value < 0.05 and log_2_fold change(FC)>|1|, based on DESeq2 results [3]. The Python (v. 3.12.2) libraries seaborn (v. 0.13.2) and matplotlib (v. 3.8.0) were used for the visualization of the results in volcano plots, Venn’s diagrams, and heatmaps. Gene ontology enrichment analysis was performed using the EnrichR website [4] with upregulated and downregulated DEGs as calculated by DESeq2.

### Human plaque collection and ex vivo culture

Human atherosclerotic plaques were collected from patients with carotid or femoral artery disease undergoing endarterectomy. Plaques were sliced into 2 mm sections, and two structurally similar pieces were placed on a moistened gelatin sponge (Spongostan Standard, Ethicon) at the medium-air interface in 6-well plates, as previously described [5]. Samples were treated with 100 nM Gal-1 or PBS in Advanced RPMI 1640 medium supplemented with antimycotic and Penicillin-Streptomycin (all Gibco) for 24 h at 37 °C, 5% CO₂. After incubation, plaques were digested in Advanced RPMI containing 1.25 mg/mL collagenase XI and 0.2 mg/mL DNase I (both Sigma-Aldrich) for 30 min at 37 °C, then passed through a 70 μm filter. Cells were stained with fixable viability dye (Invitrogen), blocked with anti-CD16/CD32 (BioLegend, 1:200), stained with surface antibodies, and then analyzed by flow cytometry.

**Major Resources table S1:** Antibodies used for immunofluorescence staining.

| **Primary antibodies** | **Dilution** | **Cat Nr** | **Company** |
| --- | --- | --- | --- |
| Rat anti-Mouse Mac-2 | 1:200 | CL8942AP | Cedarlane |
| Hamster anti-Mouse CD3e | 1:100 | 553058 | BD Bioscience |
| Rabbit anti-Mouse INOS | 1:100 | D6B6S | Cell signaling |
| Rabbit anti-Mouse Perilipin-2 | 1:200 | NB110-40877 | Novus |
| Goat anti-Mouse CD206 | 1:200 | AF2535 | R & D systems |
| **Secondary antibodies** |  |  |  |
| Donkey anti Goat IgG | 1:200 | ab96935 | Abcam |
| Dylight 488 |  |  |  |
| Donkey anti rabbit Cy3 | 1:300 | 711-165-152 | Jackson ImmunoResearch |
| Donkey anti Rat Cy5 | 1:300 | 712-175-153 | Dianova |
| sheep anti rabbit IgG | 1:200 | ab96926 | Abcam |
| Dylight 650 |  |  |  |
| Goat anti Rat IgG Dylight 488 | 1:200 | ab96887 | Abcam |
| Goat anti Hamster Cy3 | 1:300 | 127-165-160 | Jackson ImmunoResearch |

**Major Resources table S2:** Antibodies used for flow cytometry.

| Antibodies | Conjugate | Dilution | Cat Nr | Company |
| --- | --- | --- | --- | --- |
| Rat anti-Mouse CD45 | Percp cy5.5 | 1:200 | 45-0451-82 | eBioscience |
| Rat anti-Mouse CD45 | Pacific blue | 1:200 | 103125 | Biolegend |
| Rat anti-Mouse CD11b | APC | 1:200 | 101212 | Biolegend |
| Rat anti-Mouse CD11b | BV711 | 1:200 | 101242 | Biolegend |
| Rat anti-Mouse Ly6G | FITC | 1:200 | 127606 | Biolegend |
| Rat anti-Mouse Ly6G | APC cy7 | 1:200 | 127624 | Biolegend |
| Rat anti-Mouse Ly6C | AF700 | 1:200 | 128023 | Biolegend |
| Rat anti-Mouse CD115 | BV421 | 1:200 | 135513 | Biolegend |
| Rat anti-Mouse CD115 | BV605 | 1:200 | 135517 | Biolegend |
| Rat anti-Mouse F4/80 | PE | 1:200 | 123110 | Biolegend |
| Rat anti-Mouse F4/80 | PE/Cyanine7 | 1:200 | 123113 | Biolegend |
| Rat anti-Mouse B220 | PE-TR<br>(Dazzle™ 594) | 1:200 | 103258 | Biolegend |
| Rat anti-Mouse MHC-II | BV605 | 1:200 | 107639 | Biolegend |
| Rat anti-Mouse MHC-II | PE/Cyanine7 | 1:200 | 107630 | Biolegend |
| Armenian Hamster anti-Mouse CD11c | BD Horizon<br>BUV395 | 1:200 | 744180 | BD Bioscience |
| Armenian Hamster anti-Mouse CD11c | FITC | 1:200 | 117305 | Biolegend |
| Rat anti-Mouse CD3 | BV510 | 1:100 | 100234 | Biolegend |
| Rat anti-Mouse CD4 | Pacific blue | 1:200 | 100428 | Biolegend |
| Rat anti-Mouse CD8 | FITC | 1:200 | 100705 | Biolegend |
| Rat anti-Mouse CD62L | PE/Cyanine7 | 1:200 | 25-0621-82 | eBioscience |
| Rat anti-Mouse CD44 | BV605 | 1:200 | 103047 | Biolegend |
| Rat anti-Mouse CD25 | FITC | 1:200 | 102005 | Biolegend |
| Rat Anti-Mouse/Rat Foxp3 | PE | 1:100 | 72-5775-40 | eBioscience |
| Rat anti-Mouse IL-10 | PE | 1:100 | 12-7101-82 | eBioscience™ |
| Rat anti-Mouse IL-17A | APC | 1:100 | 506916 | Biolegend |
| Rat anti-Mouse IL-4 | BV421 | 1:100 | 504119 | Biolegend |
| Rat anti-Mouse IFN-γ | PE/Cyanine7 | 1:100 | 561040 | BD Biosciences |
| Armenian Hamster anti-Mouse CCR6 | PE | 1:100 | 129803 | Biolegend |
| Armenian Hamster anti-Mouse CXCR3 | PE/Cyanine7 | 1:100 | 126515 | Biolegend |
| Armenian Hamster anti-Mouse CD36 | PE | 1:200 | 102605 | Biolegend |
| Armenian Hamster anti-Mouse CD80 | APC | 1:200 | 104714 | Biolegend |
| Rat anti-Mouse CD86 | FITC | 1:200 | 105005 | Biolegend |
| Rat anti-Mouse CD206 | PE-TR<br>(Dazzle™ 594) |  | 141732 | Biolegend |
| Armenian Hamster anti-Mouse Notch1 | BV421 | 5uL/test | 130615 | Biolegend |
| Armenian Hamster anti-Mouse Notch2 | PE | 5uL/test | 130707 | Biolegend |
| Mouse anti-Human CD45 | Alexa Fluor® 700 | 5uL/test | 560566 | BD Biosciences |
| Rat anti-Mouse/Human CD11b | PE | 1:200 | 101207 | Biolegend |
| Mouse anti-Human CD68 | PE/Cyanine7 | 5uL/test | 333815 | Biolegend |
| Mouse anti-Human CD163 | Percp cy5.5 | 5uL/test | 326511 | Biolegend |
| Mouse anti-Human CD206 | BV421 | 5uL/test | 321125 | Biolegend |
| Mouse anti-Human MHC-II | BB515 | 5uL/test | 564516 | BD Biosciences |

**Major Resources table S3:**
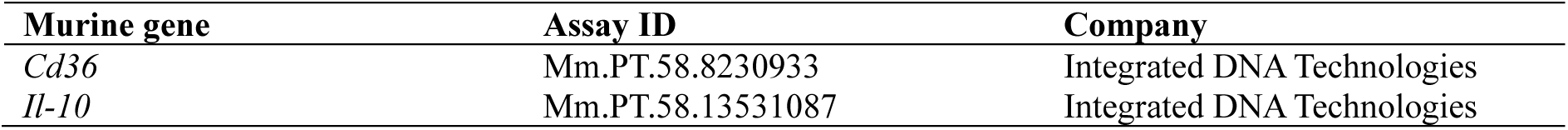
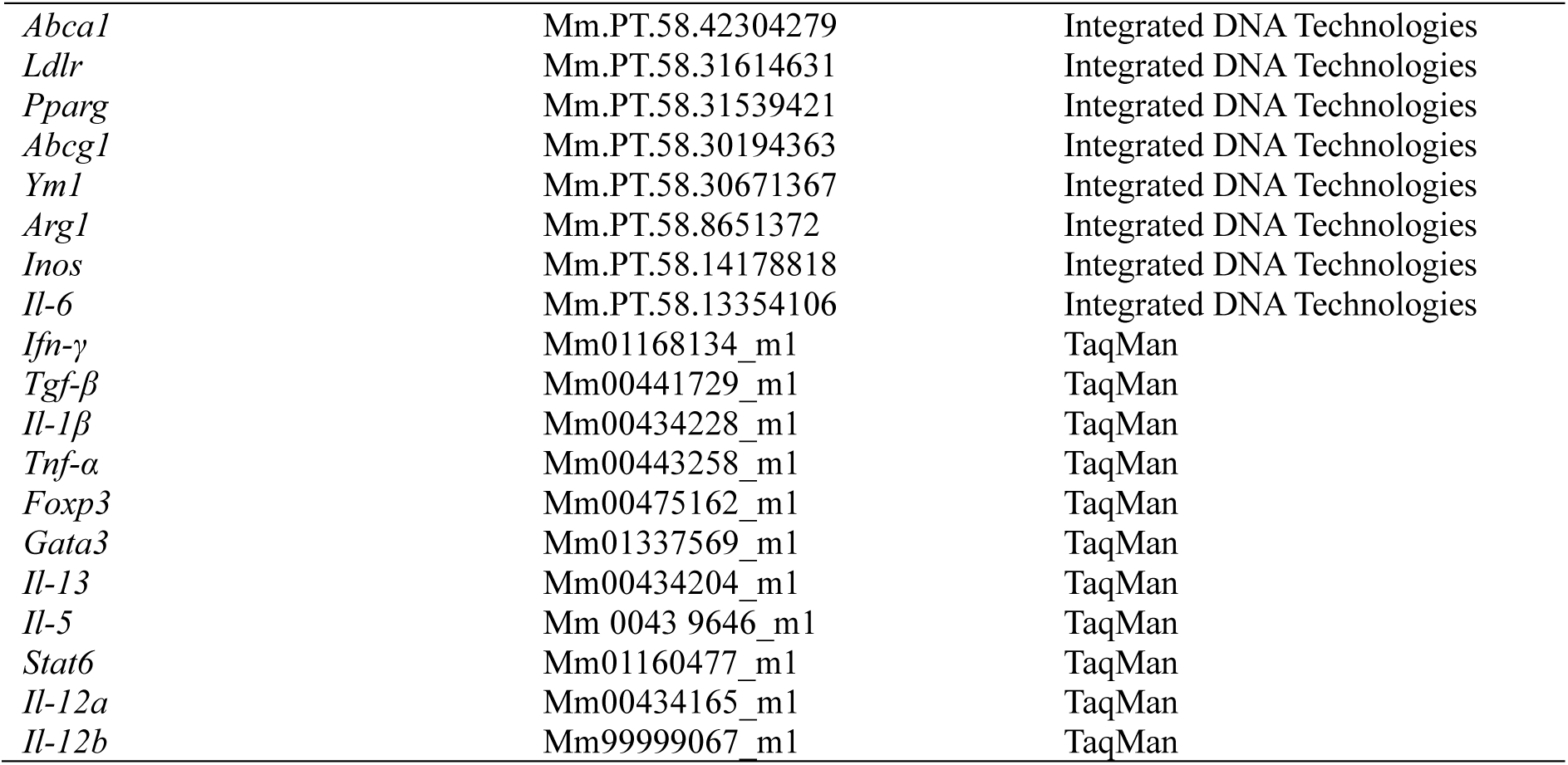
Primers used for qPCR.

**Fig. S1.**
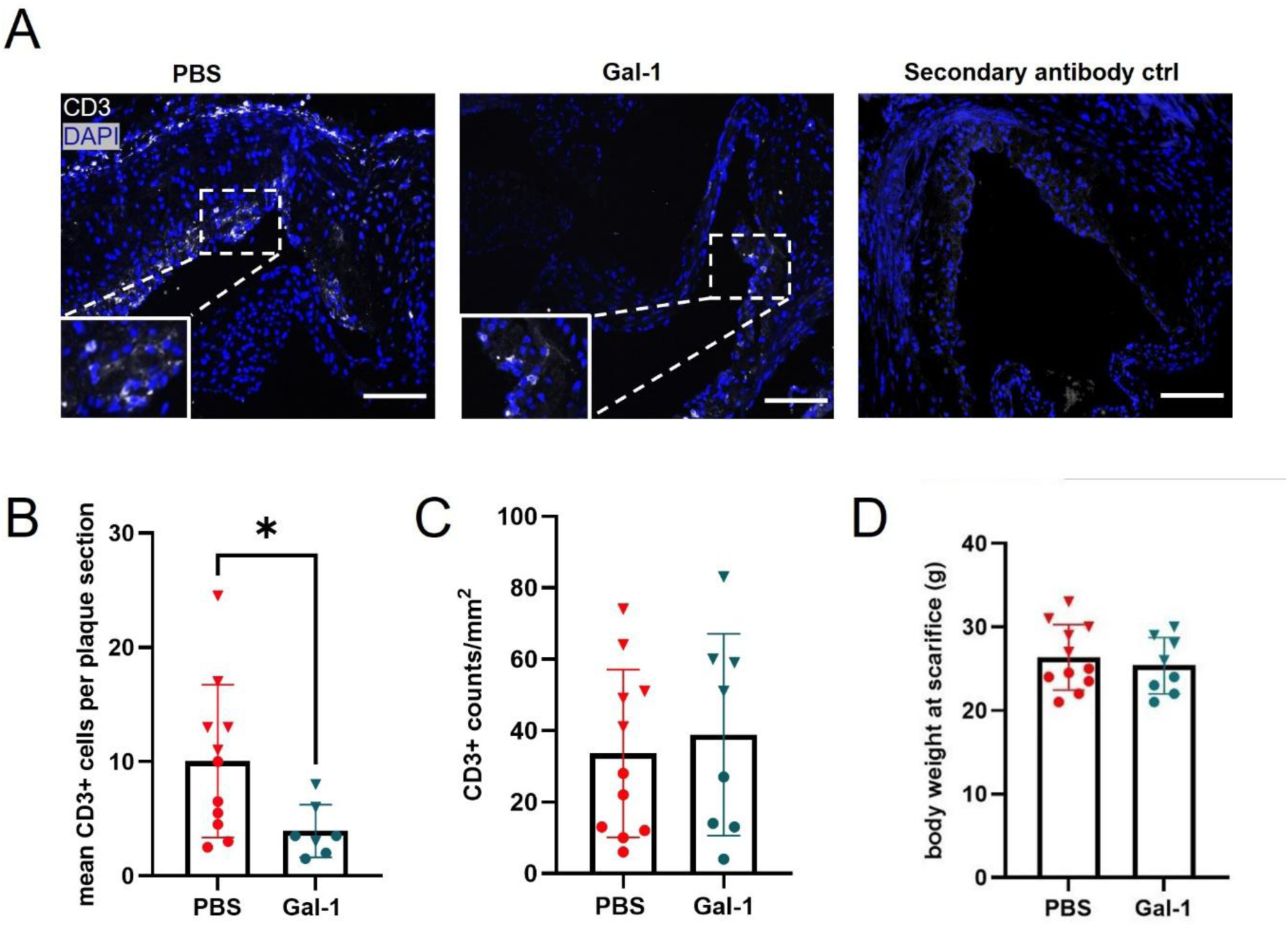
Additional data to Fig.1. **(A)** Representative images of T cell (CD3^+^) infiltration in aortic root sections from mice as shown in **Fig.1A. (B, C)** Quantification of **(B)** absolute CD3⁺ T cell counts and **(C)** CD3⁺ T cells relative to plaque area in aortic root sections. **(D)** Body weight at sacrifice after 8 weeks of WD (PBS: n=11, Gal-1: n=8). Data are presented as mean ± SEM, with individual data points shown. Bullets represent female mice, and triangles represent male mice. Scale bar: 100 μm.

**Fig. S2.**
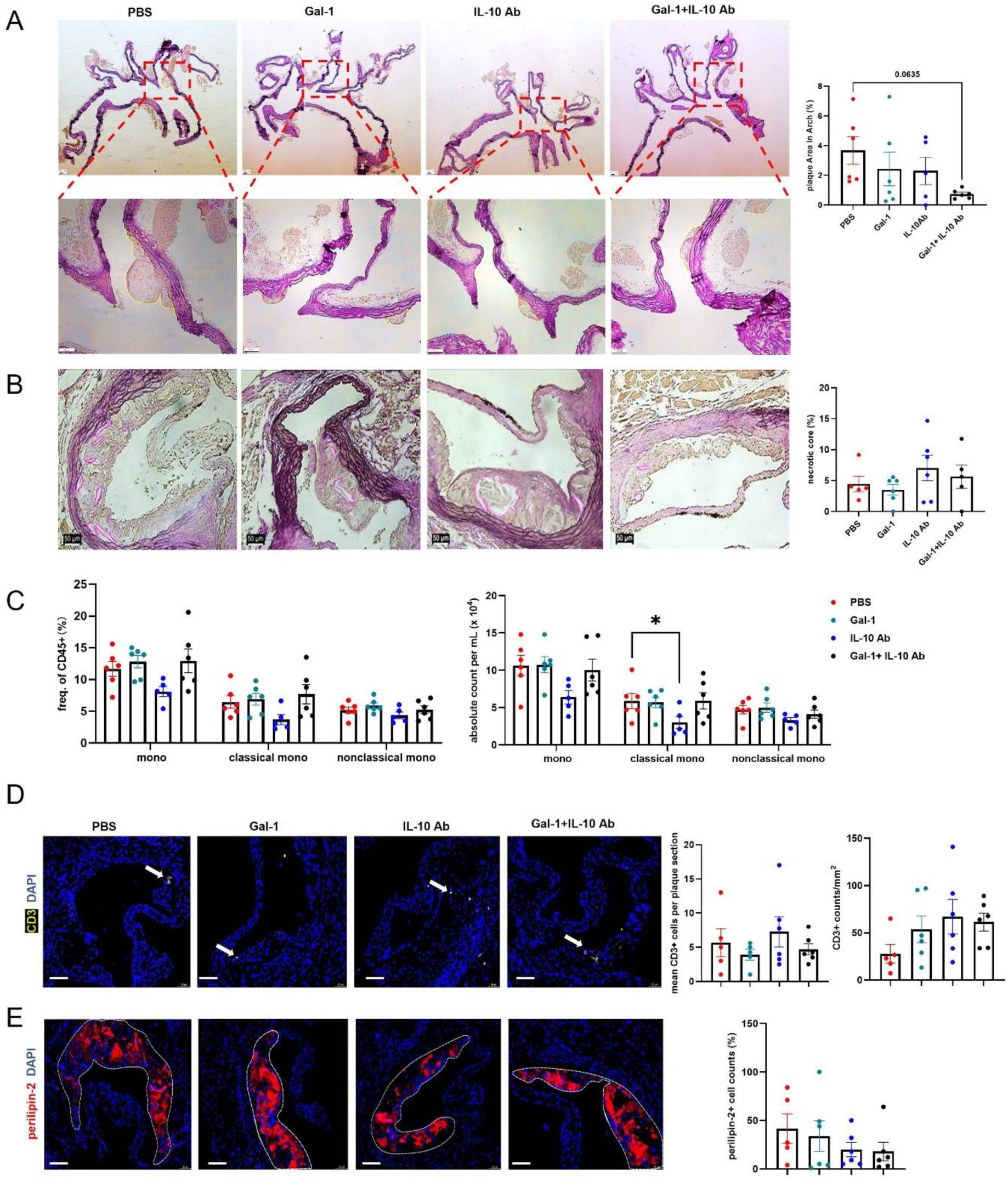
Effects of anti-IL-10 and Gal-1 treatment on aortic arch lesions. **(A)** Representative images of elastin/van Gieson (EVG) staining and quantification of plaque area in the aortic arch of *Apoe*^−/−^ female mice treated with PBS, Gal-1, IL-10 antibody (IL-10 Ab), or Gal-1 + IL-10 Ab after 4 weeks of WD (n=6 per group). **(B)** Representative images of EVG stained aortic roots showing necrotic cores and their quantification relative to plaque area (purple outline). **(C)** Percentage of monocytes (mono) and their subsets in the blood of different mouse groups following 4 weeks of WD (n=6). **(D)** Representative images and quantification of absolute T cell (CD3^+^) counts and relative to plaque area per aortic root section (PBS: n=5, other groups: n=6). **(E)** Representative images and quantification of lipid accumulation (perilipin-2^+^) per aortic root section (PBS: n=5, other groups: n=6). Data are presented as mean ± SEM. \**p* < 0.05, as determined by one-way ANOVA followed by Holm-Šídák’s post hoc test. Scale bar: 100 μm.

**Fig. S3.**
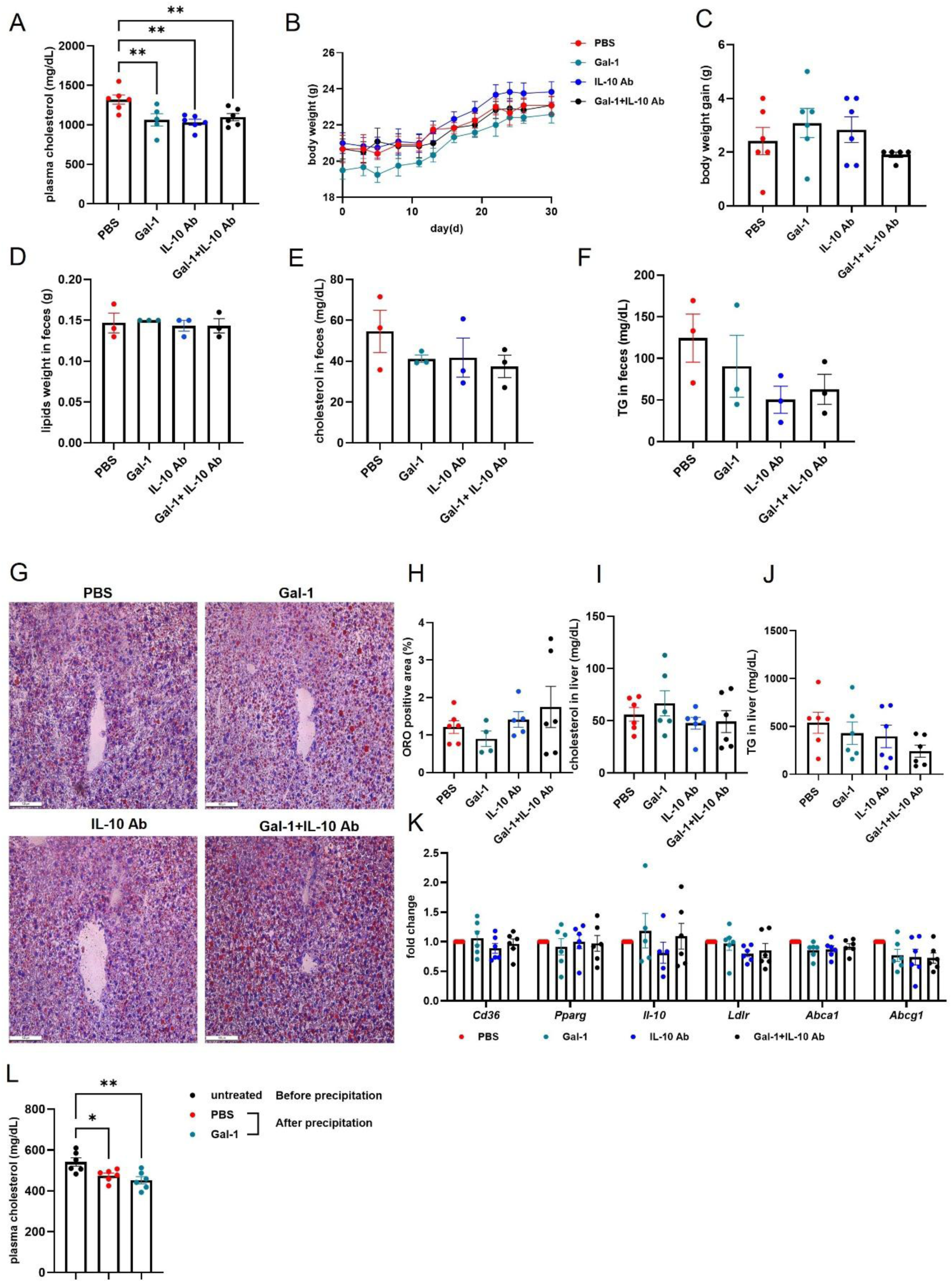
Lipid pathways after Gal-1 treatment and blocking IL-10. **(A)** Plasma total cholesterol levels were measured in *Apoe*^−/−^ mice treated as indicated after 4 weeks of WD. **(B)** Body weight and **(C)** body weight gain quantification. **(D-F)** Total lipid content, cholesterol and triglyceride levels were measured in fecal samples. **(G)** Representative ORO staining of frozen liver sections. **(H)** Quantification of ORO-positive staining in liver tissue. **(I)** Cholesterol and **(J)** triglyceride concentrations in liver tissue. **(K)** Expression of genes involved in lipid metabolism in liver tissue. **(L)** Plasma cholesterol level after Gal-1 treatment. Plasma from *Apoe*^−/−^ mice was treated with 10 µg/mL of Gal-1 or PBS for 30 minutes at room temperature. The samples were then centrifuged at 10,000 × g for 20 minutes at 4 °C to precipitate the cross-linked extracellular vesicles, and the supernatant was collected. Cholesterol levels in the supernatant were quantified post-centrifugation (n=6). Data are presented as the mean ± SEM. Statistical analysis was performed using one-way ANOVA followed by Holm-Šídák’s post hoc test. \**p* < 0.05 and \*\**p* < 0.01. Scale bar: 100 μm.

**Fig. S4.**
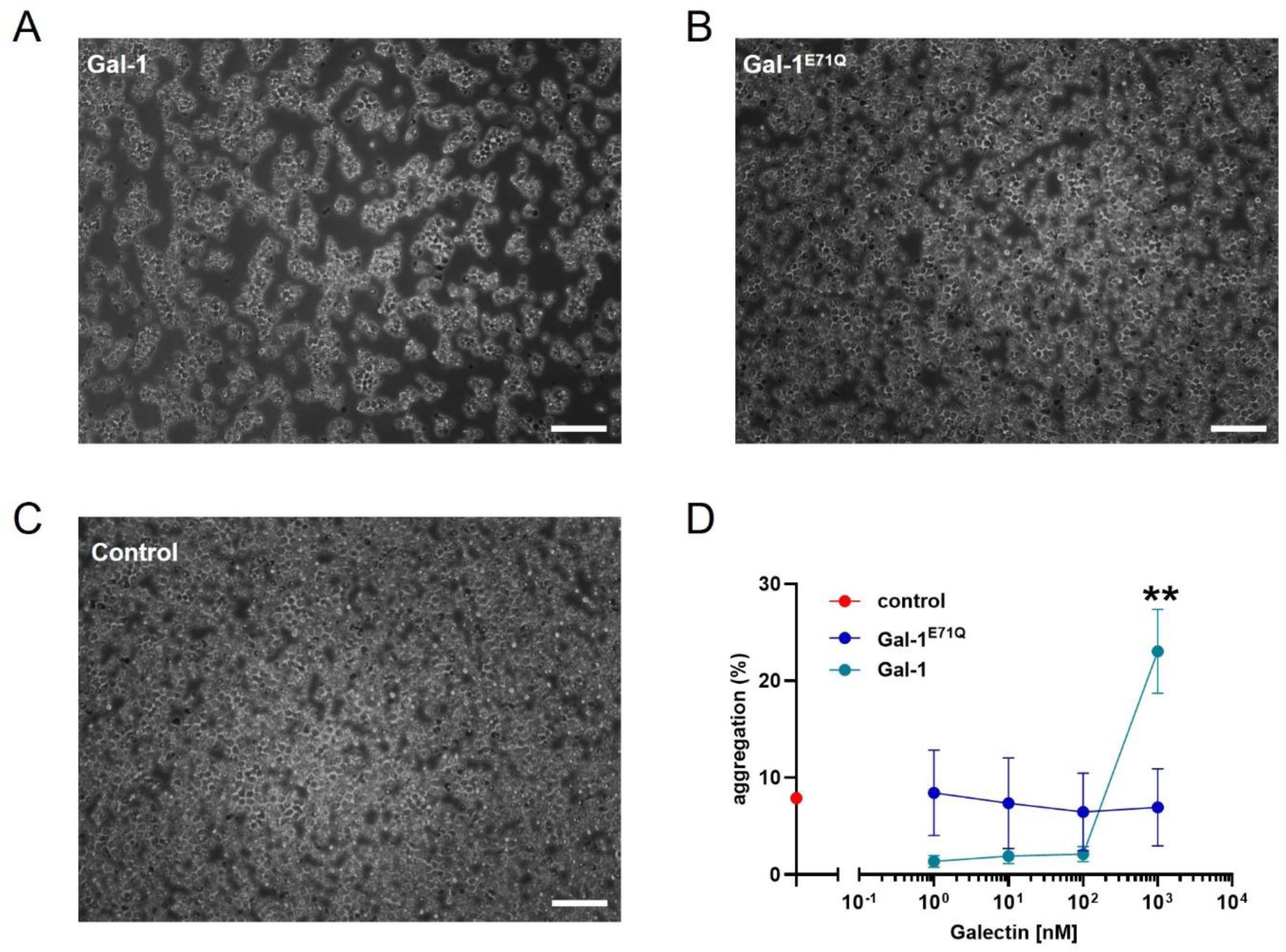
Cell aggregation induced by Gal-1 and Gal-1^E71Q^. (**A-C**) Representative images showing the aggregation of Jurkat T cells incubated with **(A)** 1 µM Gal-1, **(B)** 1 µM Gal-1^E71Q^ and **(C)** without galectin (**A-C**: n = 3). **(D)** Jurkat T cells at a concentration of 1 x 10^6^ cells/700 µL RPMI 1640 medium were incubated with increasing concentrations (from 1 nM to 1 µM) of Gal-1 (n=4) and Gal-1^E71Q^ (n=5) at 37 °C for 4 hours. The aggregation rate (doublets and clumps/all cells) was determined by flow cytometry (FSC-W vs. FSC-A). Data are presented as mean ± SEM, with statistical significance determined by two-tailed Student’s t-test. \*\**p* < 0.01. Scale bar: 100 μm.

**Fig. S5.**
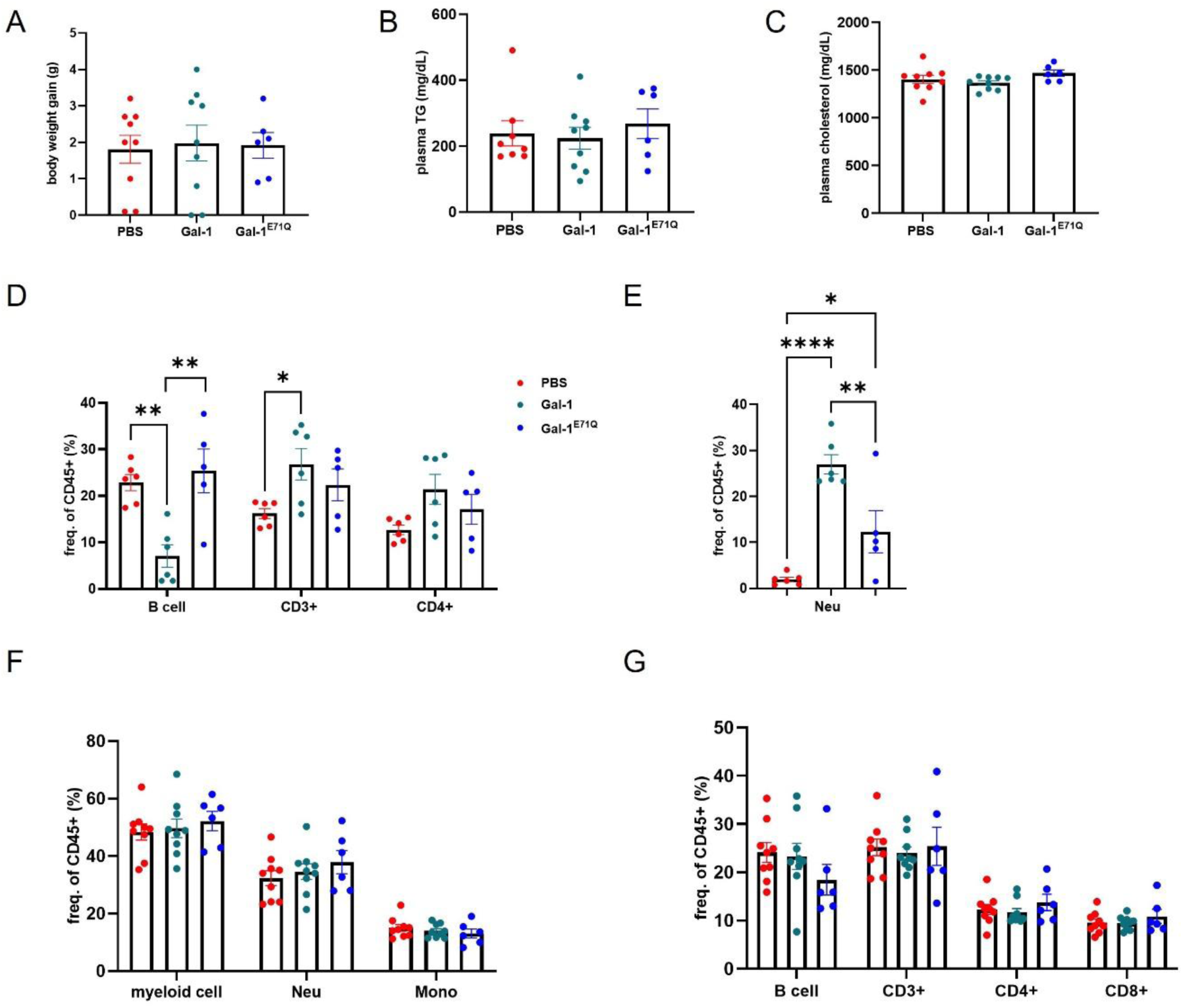
Additional data to Fig.4. **(A)** Body weight gain from mice (see scheme Fig.4A) treated with Gal-1 (n=9), Gal-1^E71Q^ (n=6) and PBS (n=9). **(B)** Plasma triglyceride levels. **(C)** Plasma cholesterol levels. **(D-G)** Percentage of leukocyte/lymphocyte subsets per CD45+ cells. (**D**) Lymphocytes in the peritoneal lavage. **(E)** Neutrophils (Neu) in the peritoneal lavage. **(F)** Myeloid cells in the blood. **(G)** Lymphocytes in the blood. \**p* < 0.05, \*\**p* < 0.01 and \*\*\*\**p* < 0.0001. Data are presented as the mean ± SEM. one-way ANOVA followed by Holm-Šídák’s post hoc test.

**Fig. S6.**
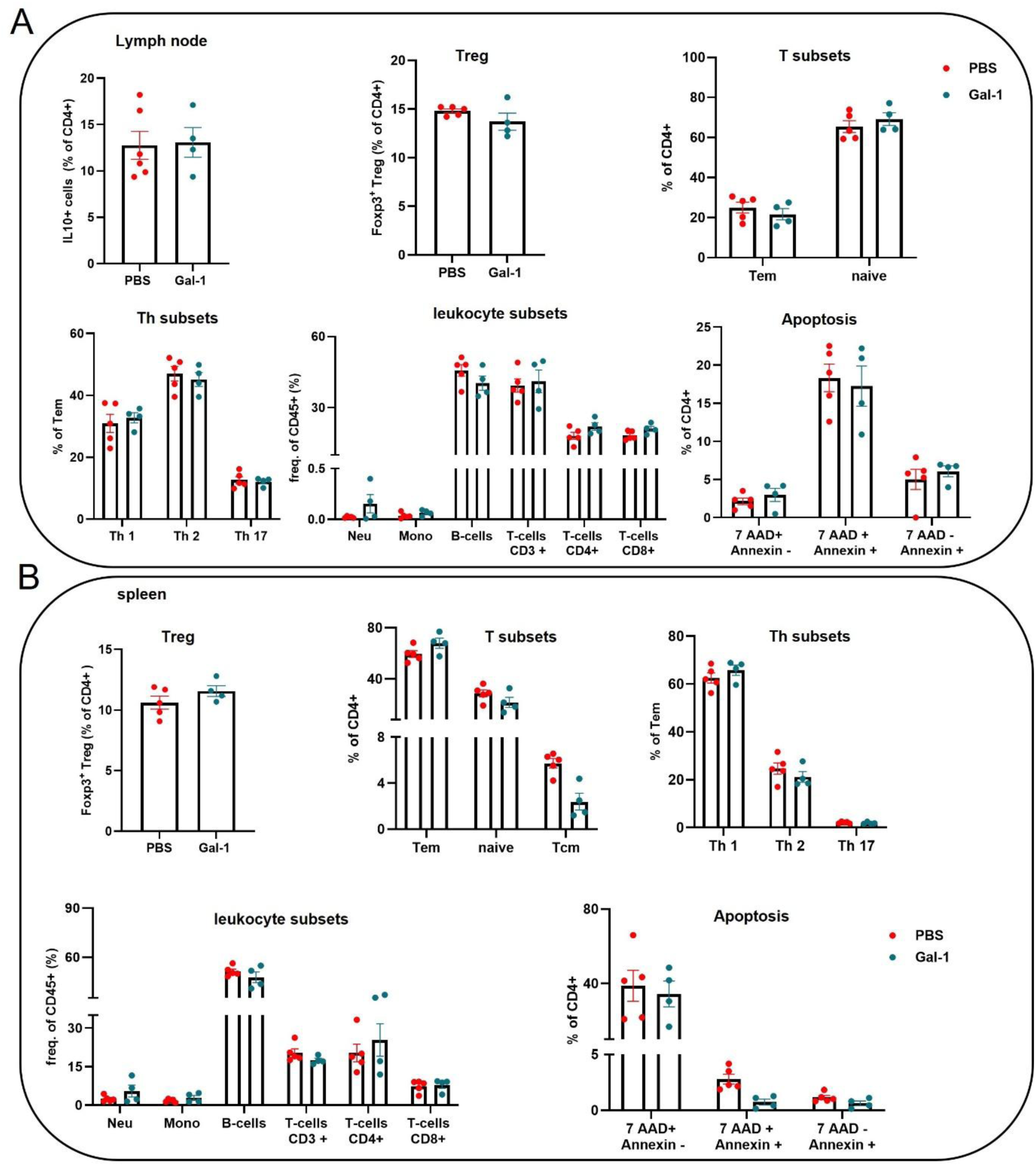
Effects of Gal-1 on leukocyte subsets and T cell viability in lymph nodes and spleen. *Apoe*^−/−^ mice were fed a WD for 8 weeks (n=4-6 per group). Frequencies of IL-10-producing T cells, Tregs (CD4^+^Foxp3^+^), effector memory (Tem: CD44^+^CD62L^-^) T cells, central memory (Tcm: CD44^+^CD62L^+^) T cells, naïve T cells (CD44^-^CD62L^+^), T helper subsets, leukocyte subsets, and CD4^+^ cell apoptosis were determined in **(A)** inguinal lymph nodes and **(B)** spleen. Data are presented as mean ± SEM. Statistical analysis was performed using a two-tailed Student’s t-test, with correction for multiple comparisons.

## References

[1] Collaborators, GBDF, Burden of disease scenarios for 204 countries and territories, 2022-2050: a forecasting analysis for the Global Burden of Disease Study 2021, Lancet 2024; 403:2204–2256.doi:10.1016/S0140-6736(24)00685-8

[2] Lacy, M, Atzler, D, Liu, R, et al., Interactions between dyslipidemia and the immune system and their relevance as putative therapeutic targets in atherosclerosis, Pharmacol Ther 2019; 193:50–62.doi:10.1016/j.pharmthera.2018.08.012

[3] Tabas, I and Lichtman, AH, Monocyte-macrophages and T cells in atherosclerosis, Immunity 2017; 47:621–634.doi:10.1016/j.immuni.2017.09.008

[4] Weber, C and von Hundelshausen, P, CANTOS trial validates the inflammatory pathogenesis of atherosclerosis: setting the stage for a new chapter in therapeutic targeting, Circ Res 2017; 121:1119–1121.doi:10.1161/CIRCRESAHA.117.311984

[5] Camby, I, Le Mercier, M, Lefranc, F, et al., Galectin-1: a small protein with major functions, Glycobiology 2006; 16:137R–157R.doi:10.1093/glycob/cwl025

[6] Cooper, D, Norling, LV and Perretti, M, Novel insights into the inhibitory effects of Galectin-1 on neutrophil recruitment under flow, J Leukoc Biol 2008; 83:1459–1466.doi:10.1189/jlb.1207831

[7] Giovannone, N, Smith, LK, Treanor, B, et al., Galectin-Glycan Interactions as Regulators of B Cell Immunity, Front Immunol 2018; 9:2839.doi:10.3389/fimmu.2018.02839

[8] Barondes, SH, Cooper, DN, Gitt, MA, et al., Galectins. Structure and function of a large family of animal lectins, J Biol Chem 1994; 269:20807–20810.doi:

[9] Eckardt, V, Miller, MC, Blanchet, X, et al., Chemokines and galectins form heterodimers to modulate inflammation, EMBO Rep 2020; 21:e47852.doi:10.15252/embr.201947852

[10] Gao, Y, Li, X, Shu, Z, et al., Nuclear galectin-1-FOXP3 interaction dampens the tumor-suppressive properties of FOXP3 in breast cancer, Cell Death Dis 2018; 9:416.doi:10.1038/s41419-018-0448-6

[11] Rostoker, R, Yaseen, H, Schif-Zuck, S, et al., Galectin-1 induces 12/15-lipoxygenase expression in murine macrophages and favors their conversion toward a pro-resolving phenotype, Prostaglandins Other Lipid Mediat 2013; 107:85–94.doi:10.1016/j.prostaglandins.2013.08.001

[12] Lightfoot, A, McGettrick, HM and Iqbal, AJ, Vascular endothelial galectins in leukocyte trafficking, Front Immunol 2021; 12:687711.doi:10.3389/fimmu.2021.687711

[13] Thiemann, S and Baum, LG, Galectins and immune responses-just how do they do those things they do?, Annual review of immunology 2016; 34:243–264.doi:10.1146/annurev-immunol-041015-055402

[14] Seropian, IM, Gonzalez, GE, Maller, SM, et al., Galectin-1 as an emerging mediator of cardiovascular inflammation: mechanisms and therapeutic opportunities, Mediators Inflamm 2018; 2018:8696543.doi:10.1155/2018/8696543

[15] Ishibashi, S, Kuroiwa, T, Sakaguchi, M, et al., Galectin-1 regulates neurogenesis in the subventricular zone and promotes functional recovery after stroke, Exp Neurol 2007; 207:302–313.doi:10.1016/j.expneurol.2007.06.024

[16] Roldan-Montero, R, Perez-Saez, JM, Cerro-Pardo, I, et al., Galectin-1 prevents pathological vascular remodeling in atherosclerosis and abdominal aortic aneurysm, Sci Adv 2022; 8:eabm7322.doi:10.1126/sciadv.abm7322

[17] Seropian, IM, Cerliani, JP, Toldo, S, et al., Galectin-1 controls cardiac inflammation and ventricular remodeling during acute myocardial infarction, Am J Pathol 2013; 182:29–40.doi:10.1016/j.ajpath.2012.09.022

[18] Perone, MJ, Bertera, S, Shufesky, WJ, et al., Suppression of autoimmune diabetes by soluble galectin-1, J Immunol 2009; 182:2641–2653.doi:10.4049/jimmunol.0800839

[19] Kopitz, J, Xiao, Q, Ludwig, AK, et al., Reaction of a programmable glycan presentation of glycodendrimersomes and cells with engineered human lectins to show the sugar functionality of the cell surface, Angew Chem Int Ed Engl 2017; 56:14677–14681.doi:10.1002/anie.201708237

[20] Janjic, A, Wange, LE, Bagnoli, JW, et al., Prime-seq, efficient and powerful bulk RNA sequencing, Genome Biol 2022; 23:88.doi:10.1186/s13059-022-02660-8

[21] Chan, IH, Van Hoof, D, Abramova, M, et al., PEGylated IL-10 activates Kupffer cells to control hypercholesterolemia, PLoS One 2016; 11:e0156229.doi:10.1371/journal.pone.0156229

[22] Shirakawa, K, Endo, J, Kataoka, M, et al., IL (Interleukin)-10-STAT3-Galectin-3 Axis Is Essential for Osteopontin-Producing Reparative Macrophage Polarization After Myocardial Infarction, Circulation 2018; 138:2021–2035.doi:10.1161/CIRCULATIONAHA.118.035047

[23] Kuhn, R, Lohler, J, Rennick, D, et al., Interleukin-10-deficient mice develop chronic enterocolitis, Cell 1993; 75:263–274.doi:10.1016/0092-8674(93)80068-p

[24] Banfer, S and Jacob, R, Galectins in intra– and extracellular vesicles, Biomolecules 2020; 10 10.3390/biom10091232

[25] Van den Bossche, J, O’Neill, LA and Menon, D, Macrophage immunometabolism: where are we (going)?, Trends Immunol 2017; 38:395–406.doi:10.1016/j.it.2017.03.001

[26] Becker, PH, Le Guillou, E, Duque, M, et al., Cholesterol accumulation induced by acetylated LDL exposure modifies the enzymatic activities of the TCA cycle without impairing the respiratory chain functionality in macrophages, Biochimie 2022; 200:87–98.doi:10.1016/j.biochi.2022.05.011

[27] Divakaruni, AS, Paradyse, A, Ferrick, DA, et al., Analysis and interpretation of microplate-based oxygen consumption and pH data, Methods Enzymol 2014; 547:309–354.doi:10.1016/B978-0-12-801415-8.00016-3

[28] Schmidt, CA, Fisher-Wellman, KH and Neufer, PD, From OCR and ECAR to energy: Perspectives on the design and interpretation of bioenergetics studies, J Biol Chem 2021; 297:101140.doi:10.1016/j.jbc.2021.101140

[29] Bisgaard, LS, Mogensen, CK, Rosendahl, A, et al., Bone marrow-derived and peritoneal macrophages have different inflammatory response to oxLDL and M1/M2 marker expression – implications for atherosclerosis research, Sci Rep 2016; 6:35234.doi:10.1038/srep35234

[30] Hirabayashi, J and Kasai, K, Effect of amino-acid substitution by site-directed mutagenesis on the carbohydrate recognition and stability of human 14-kDa β-galactoside-binding lectin, J Biol Chem 1991; 266:23648–23653.doi:

[31] Yaseen, H, Butenko, S, Polishuk-Zotkin, I, et al., Galectin-1 facilitates macrophage reprogramming and resolution of inflammation through IFN-β, Front Pharmacol 2020; 11:901.doi:10.3389/fphar.2020.00901

[32] Perillo, NL, Pace, KE, Seilhamer, JJ, et al., Apoptosis of T cells mediated by galectin-1, Nature 1995; 378:736–739.doi:10.1038/378736a0

[33] Vieceli Dalla Sega, F, Fortini, F, Aquila, G, et al., Notch signaling regulates immune responses in atherosclerosis, Front Immunol 2019; 10:1130.doi:10.3389/fimmu.2019.01130

[34] Santos, SN, Reis, C, Dias-Baruffi, M, et al., Characterization of the mechanisms underlying the crosstalk between galectins and notch in gastric cancer, BMC Proceedings 2013; 7:P65.doi:10.1186/1753-6561-7-s2-p65

[35] Parma, L, Sachs, N, Li, Z, et al., Investigating T cell recruitment in atherosclerosis using a novel human 3D tissue-culture model reveals the role of CXCL12 in intraplaque neovessels, Atherosclerosis 2024; 395 10.1016/j.atherosclerosis.2024.118385

[36] Taleb, S, Tedgui, A and Mallat, Z, IL-17 and Th17 cells in atherosclerosis: subtle and contextual roles, Arterioscler Thromb Vasc Biol 2015; 35:258–264.doi:10.1161/ATVBAHA.114.303567

[37] Bollyky, PL, Wu, RP, Falk, BA, et al., ECM components guide IL-10 producing regulatory T-cell (TR1) induction from effector memory T-cell precursors, Proc Natl Acad Sci U S A 2011; 108:7938–7943.doi:10.1073/pnas.1017360108

[38] Ilarregui, JM, Croci, DO, Bianco, GA, et al., Tolerogenic signals delivered by dendritic cells to T cells through a galectin-1-driven immunoregulatory circuit involving interleukin 27 and interleukin 10, Nat Immunol 2009; 10:981–991.doi:10.1038/ni.1772

[39] Shi, H, Guo, J, Yu, Q, et al., CRISPR/Cas9 based blockade of IL-10 signaling impairs lipid and tissue homeostasis to accelerate atherosclerosis, Front Immunol 2022; 13:999470.doi:10.3389/fimmu.2022.999470

[40] Stoger, JL, Boshuizen, MC, Brufau, G, et al., Deleting myeloid IL-10 receptor signalling attenuates atherosclerosis in LDLR−/−mice by altering intestinal cholesterol fluxes, Thromb Haemost 2016; 116:565–577.doi:10.1160/TH16-01-0043

[41] Baek, JH, Kim, DH, Lee, J, et al., Galectin-1 accelerates high-fat diet-induced obesity by activation of peroxisome proliferator-activated receptor gamma (PPARγ) in mice, Cell Death Dis 2021; 12:66.doi:10.1038/s41419-020-03367-z

[42] Chen, Y, Yang, M, Huang, W, et al., Mitochondrial metabolic reprogramming by CD36 signaling drives macrophage inflammatory responses, Circ Res 2019; 125:1087–1102.doi:10.1161/CIRCRESAHA.119.315833

[43] Guda, MR, Tsung, AJ, Asuthkar, S, et al., Galectin-1 activates carbonic anhydrase IX and modulates glioma metabolism, Cell Death Dis 2022; 13:574.doi:10.1038/s41419-022-05024-z

[44] Starossom, SC, Mascanfroni, ID, Imitola, J, et al., Galectin-1 deactivates classically activated microglia and protects from inflammation-induced neurodegeneration, Immunity 2012; 37:249–263.doi:10.1016/j.immuni.2012.05.023

[45] Garin, MI, Chu, CC, Golshayan, D, et al., Galectin-1: a key effector of regulation mediated by CD4+CD25+ T cells, Blood 2007; 109:2058–2065.doi:10.1182/blood-2006-04-016451

[46] Mor, A, Planer, D, Luboshits, G, et al., Role of naturally occurring CD4+ CD25+ regulatory T cells in experimental atherosclerosis, Arterioscler Thromb Vasc Biol 2007; 27:893–900.doi:10.1161/01.ATV.0000259365.31469.89

[47] Harrington, LE, Hatton, RD, Mangan, PR, et al., Interleukin 17-producing CD4+ effector T cells develop via a lineage distinct from the T helper type 1 and 2 lineages, Nat Immunol 2005; 6:1123–1132.doi:10.1038/ni1254

[48] Perone, MJ, Larregina, AT, Shufesky, WJ, et al., Transgenic galectin-1 induces maturation of dendritic cells that elicit contrasting responses in naive and activated T cells, J Immunol 2006; 176:7207–7220.doi:10.4049/jimmunol.176.12.7207

[49] de la Fuente, H, Cruz-Adalia, A, Martinez Del Hoyo, G, et al., The leukocyte activation receptor CD69 controls T cell differentiation through its interaction with galectin-1, Mol Cell Biol 2014; 34:2479–2487.doi:10.1128/MCB.00348-14

[50] McGeachy, MJ, Bak-Jensen, KS, Chen, Y, et al., TGF-β and IL-6 drive the production of IL-17 and IL-10 by T cells and restrain T(H)-17 cell-mediated pathology, Nat Immunol 2007; 8:1390–1397.doi:10.1038/ni1539

[51] Mills, KHG, IL-17 and IL-17-producing cells in protection versus pathology, Nat Rev Immunol 2023; 23:38–54.doi:10.1038/s41577-022-00746-9

[52] Gagliani, N, Vesely, MCA, Iseppon, A, et al., Th17 cells transdifferentiate into regulatory T cells during resolution of inflammation, Nature 2015; 523:221–225.doi:10.1038/nature14452

## Supplementary References

[1] Kopitz J, Xiao Q, Ludwig AK, Romero A, Michalak M, et al. Reaction of a programmable glycan presentation of glycodendrimersomes and cells with engineered human lectins to show the sugar functionality of the cell surface. Angew Chem Int Ed Engl 2017; 56:14677–14681.doi:10.1002/anie.201708237

[2] Janjic A, Wange LE, Bagnoli JW, Geuder J, Nguyen P, et al. Prime-seq, efficient and powerful bulk rna sequencing. Genome biology 2022; 23:88.doi:10.1186/s13059-022-02660-8

[3] Love MI, Huber W and Anders S. Moderated estimation of fold change and dispersion for rna-seq data with deseq2. Genome biology 2014; 15:550.doi:10.1186/s13059-014-0550-8

[4] Xie Z, Bailey A, Kuleshov MV, Clarke DJB, Evangelista JE, et al. Gene set knowledge discovery with enrichr. Curr Protoc 2021; 1:e90.doi:10.1002/cpz1.90

[5] Parma L, Sachs N, Li Z, Merchant K, Sobczak N, et al. Investigating t cell recruitment in atherosclerosis using a novel human 3d tissue-culture model reveals the role of cxcl12 in intraplaque neovessels. Atherosclerosis 2024; 395. doi:10.1016/j.atherosclerosis.2024.118385

